# Spatial Regulation of Lck Activation at the CD8 Immune Synapse Revealed by a FRET-Based Biosensor

**DOI:** 10.1101/2025.10.24.684336

**Authors:** Clara Meana, Gonzalo San-José, María A. Balboa, Javier Casas

**Affiliations:** Unidad de Excelencia Instituto de Biomedicina y Genética Molecular (IBGM), Consejo Superior de Investigaciones Científicas (CSIC) - Universidad de Valladolid (UVa), Spain; Departamento de Bioquímica y Biología Molecular y Fisiología, Facultad de Medicina, Universidad de Valladolid, 47003 Valladolid, Spain; Departamento de Pediatría e Inmunología, Universidad de Valladolid, Valladolid 47005, Spain; Lipid Metabolism and Inflammation Group, IBGM, CSIC-UVA, 47003 Valladolid, Spain; Centro de Investigación Biomédica en Red de Diabetes y Enfermedades Metabólicas Asociadas (CIBERDEM), 28029 Madrid, Spain

**Keywords:** CD8, TCR, LCK, Immune Synapse, FRET, Spatial Regulation

## Abstract

T cell receptor (TCR) signaling is critically dependent on the Src-family kinase Lck, whose activation is tightly regulated both spatially and conformationally during antigen recognition. Here, we employ an improved second-generation FRET-based biosensor (TqLckV2.3) to visualize Lck conformational dynamics in live T cells with high spatial resolution. Upon TCR engagement, we observe a paradoxical increase in whole-cell FRET signal, which immunolabeling reveals to be due to selective internalization of inactive Lck (pY^505^), while active Lck (pY^394^) remains membrane-associated and enriched at the immune synapse (IS). Using CD8α mutants that disrupt Lck binding, we demonstrate that free Lck undergoes more pronounced conformational activation than CD8-bound Lck. Furthermore, we show that C-terminal Src kinase (Csk) preferentially phosphorylates free Lck at Y^505^, while CD45 suppresses its activation via dephosphorylation of Y^394^, suggesting a dual regulatory mechanism that maintains free Lck in an inactive state under resting conditions. High-resolution imaging confirms sustained activation of Lck at the IS and transient inactivation at the periphery, revealing a spatially confined signaling architecture. These findings uncover a novel regulatory mechanism involving selective internalization and spatial segregation of Lck conformations and establish TqLckV2.3 as a powerful tool for dissecting TCR signaling dynamics.

## INTRODUCTION

T cell activation is a highly regulated process initiated by the engagement of the T cell receptor (TCR) with peptide–major histocompatibility complex (pMHC) molecules on antigen-presenting cells (APCs). This interaction triggers a cascade of intracellular signaling events, many of which are orchestrated by the Src-family kinase Lck. Lck plays a pivotal role in phosphorylating immunoreceptor tyrosine-based activation motifs (ITAMs) within the CD3 complex, thereby initiating downstream signaling pathways essential for T cell function and fate decisions [1].

Despite its central role, the mechanisms governing Lck activation and spatial regulation during TCR engagement remain incompletely understood. Lck exists in multiple conformational states, regulated by phosphorylation at two key tyrosine residues: Y^394^ (activating) and Y^505^ (inhibitory) [2, 3]. The dynamic balance between these states is influenced by interactions with coreceptors (CD4/CD8), phosphatases, and kinases, as well as by the spatial organization of signaling complexes at the immune synapse [4, 5].

Our previous work has proposed that free Lck, not bound to CD8, may be primarily responsible for initiating TCR signaling, while CD8-bound Lck contributes to signal amplification or stabilization [6, 7]. However, direct evidence supporting this model has been limited by the lack of tools capable of resolving Lck conformational changes in live cells with high spatial and temporal resolution.

The lymphocyte-specific protein tyrosine kinase Lck plays a critical role in T cell receptor (TCR) signaling initiation, yet the steady-state level of constitutively active Lck in resting T cells remains one of the most contentious issues in T cell biology. Studies have reported dramatically conflicting values, ranging from as low as ∼2% to as high as ∼40% of the total Lck pool existing in a constitutively active state—a striking 20-fold discrepancy with profound implications for our understanding of TCR signaling thresholds and T cell activation mechanisms. [8, 9]. This controversy has been largely attributed to methodological artifacts [9].Despite these quantitative uncertainties, the functional significance of constitutive Lck activity is well-established [8, 10]. The resolution of this controversy requires not only improved methodological approaches that control for post-lysis artifacts, but also a deeper understanding of how different detection methods may be measuring distinct aspects of Lck activation, including conformational states versus phosphorylation-dependent enzymatic activity [11, 12].

We used an improved second-generation FRET-based biosensor, TqLckV2.3, optimized for monitoring Lck conformational dynamics in T cells. Ratiometric FRET enables real-time monitoring of dynamic molecular interactions with high temporal resolution, which is critical for capturing the fast kinetics of early T cell signaling events. Unlike Fluorescence Lifetime Imaging Microscopy (FLIM), which requires longer acquisition times and complex data fitting to resolve fluorescence lifetimes, ratiometric FRET can be implemented with standard widefield or confocal microscopy, allowing for faster image acquisition and analysis. This speed is essential for studying transient signaling intermediates and spatially localized activation patterns at the immunological synapse. TqLckV2.3 enables the visualization of Lck conformational states in live T cells and provides insights into the spatial compartmentalization of active and inactive Lck during immune synapse formation. In this study, we combine live-cell imaging, phospho-specific labeling, and mutant T cell models to dissect the conformational regulation of Lck during antigen recognition. Our findings reveal a novel mechanism of Lck regulation involving the selective internalization of its inactive form and demonstrate that free Lck undergoes more substantial conformational changes than CD8-bound Lck upon TCR engagement

## MATERIALS AND METHODS

### Constructs and Cells

CHO cells expressing the tetracycline repressor (Thermo, TRex) under blasticidin selection (10 μg/ml) were grown in Ham’s F12 media with 10% (vol/vol) FCS, 100 U/ml of penicillin, 10 mg/ml of streptomycin, hygromycin 0.3 mg/ml and G418 0.8 mg/ml. When 50 ng/ml tetracycline/doxycycline (DOX) is added, it binds the Tet repressor, causing it to release from the operator sites, allowing transcription of the target genes. Hybridomas expressing the OT-I TCR with or without CD8α and CD8β were made by retroviral transduction of 58α-β cells [13]. In brief, this is a variant of the DO-11.10.7 mouse T cell hybridoma which does not express functional T cell receptor α/β chains nor CD8 chains, but it does express OT-I α and β-chains after retroviral transduction and is referred as OVAαβ cells (Fig S1). From this line, we created two variants: (i) OT-I WT cells expressing CD8α and CD8β (Fig S1, 17αβ), and (ii) OT-I CxCP cells (Fig S1 CxCP), expressing CD8α-CxCP mutant together with CD8β, which prevents Lck association through the CXCP motif [14]. All OT-I hybridoma cells were maintained in IMDM medium containing 10% fetal bovine serum (FBS), 2 mM L-glutamine, 100 U/ml penicillin/streptomycin, 50 μM β-mercaptoethanol, 500 μg/ml G418 (selecting for TCRα) and 3 μg/ml puromycin (selecting for TCRβ) and FACS sorted for CD8 expression when necessary.

The plasmid pEF-TqLckV2.3 was provided by Dr. Burkhart Schraven from Magdeburg University. TqLckV2.3 was amplified by PCR adding 5’Hind-III and Not-I 3’specific restriction sites and inserted into retroviral vector pBMN-Z (S. Kinoshita and G. Nolan, www.stanford.edu/group/nolan). All the mutants were made using the Quickchange Site Mutagenesis Kit (Stratagene). pHR-Csk-P2A-Cbp and pHCM-mCherry-CD45_cyt_CD43_ex_ were kindly provided by Dr. John James from Oxford University.

### Surface staining and Flow cytometry

OT-I hybridoma cells were stained with the following conjugated antibodies BV450 anti-CD8α (eBioscience, clone 53-6.7), APC anti-CD8β (eBioscience, clone H35-17.2) or PE/Cy7 anti-CD45 (BioLegend, clone 30-F11). All flow cytometry was conducted on a 5-laser spectral cytometer (Aurora, Cytek) at the Institute of Biomedicine and Molecular Genetics flow cytometry facility. Flow cytometry data were analyzed in FlowJo (BD Biosciences) and exported where necessary.

### Lentiviral and Retroviral Transduction

VSV-G–pseudotyped lentiviral particles were produced by co-transfection of 293FT cells with transfer constructs (CD45_cyt_-CD43_ex_ and Csk-P2A-Cbp) and the compatible packaging plasmids pMD2.G and psPAX2 in the presence of PEI 25K. Ecotropic retroviral particles were produced by co-transfection of Plat-E cells with transfer constructs (pBMN-TqLckV2.3, pBMN-CD8α-CxCP or pBMN-CD8β) and the compatible packaging plasmid pCL-Eco in the presence of PEI 25K. Viruses were harvested at 48 and 72 hours after transfection. Viral transduction of OT-I T cell hybridomas were carried out using concentrated viral particles with AMICON Ultracel 100K concentrating tubes (Millipore) in the presence of 8 μg/ml Polybrene (Sigma-Aldrich), and infected cells were selected by FACS sorting or antibiotic resistance and subjected to imaging experiments.

### TCR crosslinking stimulation

For soluble antibody–based stimulation, OT-I T cell hybridomas were incubated with biotinylated anti-CD3ε (BioLegend, clone 145-2C11) and biotinylated anti-CD8α (BioLegend, clone 53-6.7) at a final concentration of 2.5 µg/ml each for 20 min at 4 °C in FACS staining solution (HBSS supplemented with 1%BSA and 5% heat inactivated human serum). After washing at 4°C and centrifugation, antibody cross-linking was induced by addition of streptavidin (1 µg/ml) at 37° C for the indicated times or at the indicated time point during live-cell FRET acquisition. Stimulation was stopped by adding 1 volume of 8% paraformaldehyde (PFA). This procedure generated a synchronized and global activation of the TCR signaling machinery, as previously described for CD3/CD8 co-cross-linking. When required, cells were pre-treated with the Src-family kinase inhibitor PP2 (10 µM, 20 min at 37 °C) before antibody addition to prevent Lck activation and to establish inhibitory controls for FRET measurements or phospho-specific analyses. In all cases, cells were maintained in HBSS supplemented with 10 mM HEPES, 1mM CaCl_2_ and 1 mM MgCl_2_ buffer during stimulation, and untreated samples received vehicle alone.

### Conjugate Assay

1.0×10^5^ OT-I hybridomas and 1.0×10^5^ DOX induced and Cy5 labeled CHO cells were added to 96-well round bottom wells in a total volume of 50 ml and incubated for the indicated times at 37°C. The cells were pipetted up and down three times after the incubation period before fixing in 4% PFA at indicated time points. After fixing for 20 min at RT, the cells were washed in PBS, and the PFA was inactivated by 10 mM Tris (pH 7.4) in PBS for 5 min at RT. The cell conjugates were examined by flow cytometry based on simultaneous expression of mVenus (OT-I hybridoma) and Cy5 (Cy5-labeled CHO cell). Conjugate frequency was calculated as the percentage of double-positive T cell–APC events among the OT-I (mVenus positive) population.

### Fixed Cell Confocal FRET Imaging

OT-I hybridoma cells expressing the TqLckV2.3 biosensor were added onto poly-L-lysine–coated glass coverslips containing a pre-plated monolayer of DOX-induced CHO cells expressing scH-2Kᵇ-OVA and surface-labeled with Cy5. After 15 min to allow conjugate formation, non-interacting T cells were removed by gentle PBS washes and samples were fixed in 4% PFA for 20 min at room temperature. Cells were imaged on a Leica confocal system TCS SP5X, with a HCS Plan Apo CS 63X/1.4 NA oil immersion lens, correct localization of the sensor was confirmed by mVenus imaging. Cy5 fluorescence was obtained by 647 nm excitation and 657-700 nm emission. For ratio imaging, Turqouise (Tq) fluorescence was obtained by 458 nm excitation and 465-510 nm emission, mVenus by 515 nm excitation and 525-580 nm emission and uncorrected FRET by 458 nm excitation and 525-580 nm emission. Tq and FRET images where simultaneously acquired while Venus was acquired alternately. ImageJ /FIJI software with customized plugins was used to correct the background from raw images and to create ratio images and export raw data to Microsoft Excel for further ratio and correction calculations.

For whole-cell analyses, z-stacks encompassing the entire T cell (from the basal contact plane to the apical membrane) were Z-projected (sum-intensity) in ImageJ/Fiji to generate 2D maps representing the entire cell volume. When required for presentation confocal z-stacks images were deconvolved using an ImageJ parallel iterative plugin (Wiener Filter Preconditioned Landweber (WPL) method) after calculating the experimental setup point spread function (PSF) using sub resolution (0.17 μm) fluorescent beads.

### Live Cell FRET Imaging

A dual superfast external excitation/emission wheels specifically designed for live FRET were used for imaging, allowing almost simultaneous acquisition of donor and acceptor emission during donor excitation (20 ms delay). This consisted in a Leica K5 sCMOS camera attached to a Leica Dmi8 inverted microscope equipped with aforementioned ultrafast external filter wheels and a CoolLED 3 illumination system. Cells expressing Lck activity FRET-based sensor (TqLckV2.3) were cultured in glass-bottomed culture dishes (Mat-Tek). When improved adhesion was required, the glass was coated with poly-L-lysine (Sigma). The culture medium was replaced with HBSS supplemented with 10 mM HEPES, pH 7.4, 1.3 mM CaCl₂, and 1.3 mM MgCl₂. Cells were visualized at 37°C using an HC PL FLUOTAR 63X/1.3 NA oil immersion lens and an infrared focus maintenance system. Proper localization of the sensor was confirmed by obtaining mVenus images. For ratiometric imaging, Tq fluorescence was obtained using an excitation filter at 430/24 nm and an emission filter at 470/24 nm, Venus using an excitation filter at 500/20 nm and an emission filter at 535/30 nm, and FRET, without correction, using the Tq excitation filter at 430/24 nm and the Venus emission filter. The acquisition of the three channels was virtually simultaneous. We recorded a 2 min baseline before adding T cells acquiring images each 5 s. The selection of the IS and other dynamic regions in live-cell FRET imaging must be performed manually due to the dynamic nature of conjugate formation and antigen recognition. The initial contact point is clearly identifiable and marked as time zero. From that point onward, regions of interest (ROIs) were carefully and manually drawn for each subsequent time frame, considering the plasma membrane thickness as indicated by the mVenus channel. ImageJ/FIJI software with custom plugins was used to correct both the movement and the background of the raw images and to create pixel-by-pixel FRET/Tq ratio images.

### Immunocytochemistry Analyses

For phospho-specific staining of Lck, cells were fixed with 4% PFA for 20 min in suspension and then permeabilized with 0.1% Triton X-100 for 20 min at room temperature. Cells were treated with 5% goat serum for 30 min and incubated with antibodies against 125 ng/ml of phospho Y^505^, phospho Y^394^, total Lck (Cell Signaling, clone 73A5) or 100 ng/ml phospho-Tyr (4G10 clone, Merck) for 1h at RT. After washing, the cells were incubated with 1 μg/ml spectrally appropriate Alexa Fluor-conjugated Fab fragments against primary species antibodies (Thermo) for 1h at RT. After intracellular staining, nuclei were counterstained with the DNA binding dye Hoechst 33342 (Thermo). All images were captured with a Leica confocal system TCS SP5X inverted microscope with a HCS Plan Apo CS 63X/1.4 NA oil immersion lens. Leica Application Suite Advanced Fluorescence software was used for the capture, and ImageJ/FIJI was used for deconvolution (Parallel Iterative 3D plugin), image presentation and quantification.

### Immunoblots

Cells were lysed in a buffer 1% Triton X-100, 50 mM Tris, pH 7.4, 150 mM NaCl, 1 mM EDTA, 1 mM EGTA, 5 mM Na_4_P_2_O_7_, 50 mM sodium β-glycerol phosphate, 270 mM sucrose, 0.1% β-mercaptoethanol, 1 mM Na_3_VO_4_, 10 mM NaF, 1 mM phenylmethyl sulphonyl fluoride and a protease inhibitor ‘cocktail’ (Sigma). Cell proteins were separated by 10% SDS–polyacrylamide gel electrophoresis and transferred to nitrocellulose membranes. The membranes were incubated with anti-Csk (BD Biosciences, clone 52/Csk) or anti-β-actin (Sigma-Aldrich, clone AC15) followed by incubation with goat anti-mouse IgG IR-Dye 800CW-conjugated (LiCor) and detected and accurately quantified with a LiCor Odyssey Fc infrared imaging system.

### Statistical Analysis

Statistical details of experiments are indicated in the figure legends. All data analyses were performed with Prism software (GraphPad). No statistical analysis was used to predetermine sample size. Data are represented as mean ± SEM. Statistical significance was determined by one-way ANOVA with Tukey’s post hoc test or unpaired two-tailed Student’s *t* test. *p* < 0.05 was considered statistically significant. Some data required multiple t-test, with a false discovery rate of 1% based on two stage step up by Benjamini, Krieger and Yekutieli.

## RESULTS

### Functionality of an improved second-generation FRET-based Lck biosensor

To monitor Lck conformational dynamics in live cells, we employed an optimized second-generation FRET-based biosensor, TqLckV2.3. In this construct, the original CFP and YFP fluorophores were replaced with monomeric Turquoise2 and monomeric Venus, respectively as in the second generation sensor [11, 15, 16], although the entire insert was codon-optimized to minimize recombination events during retroviral cloning. As in previous designs, the donor fluorophore Tq was repositioned between the SH3 and SH2 domains of Lck, based on prior findings demonstrating that this configuration more accurately recapitulates TCR-mediated signaling (Fig. 1a) [11, 15, 17]. Importantly, cloning the biosensor into a retroviral vector enables efficient transduction of hard-to-transfect cells such as T cells. This approach generates stable and uniformly expressing cell populations (Fig. S2), thereby reducing variability compared to transient transfection. Such uniformity is essential for obtaining reliable FRET measurements in functional assays.

**Figure 1.**
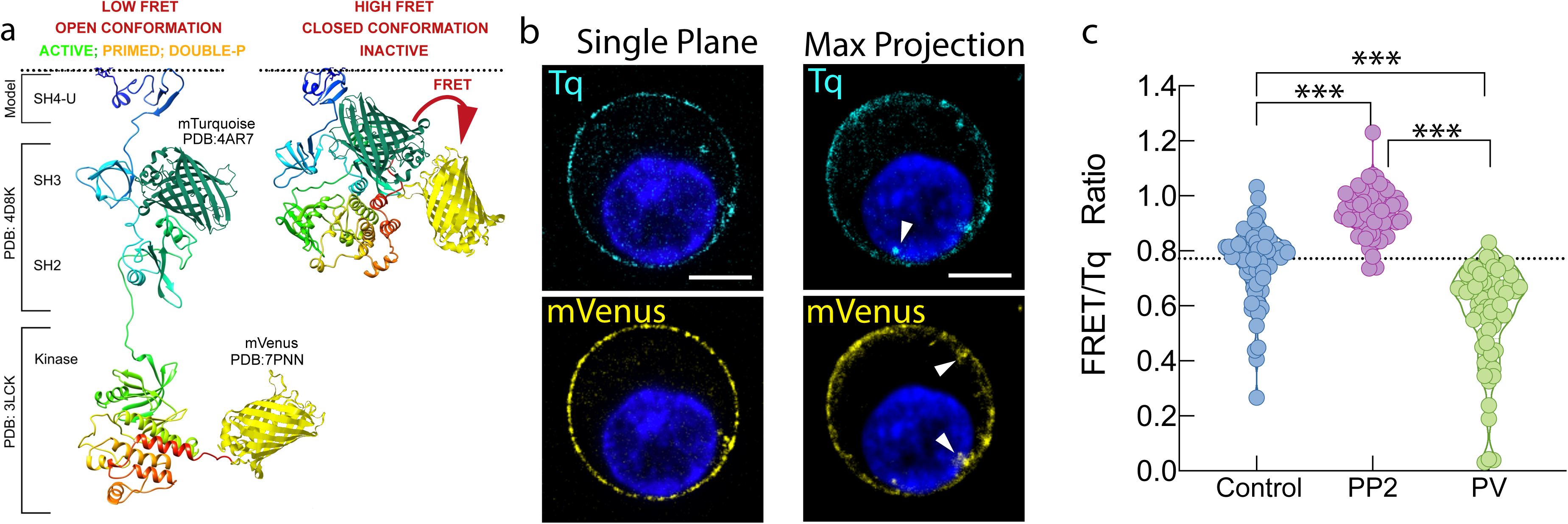
Schematic representation of the conformational changes that result in FRET changes in the TqLckV2.3 biosensor. a) The model of the open conformation (left pane) and closed conformation (right pane), modified from Prakaash et al. Sci Rep (2022). b) OT-I WT T cell hybridoma expressing TqLckV2.3 biosensor were analyzed by confocal microscopy and a single plane, or a Z max projection is shown. Arrows indicate endosomal pool of Lck. Scale bar, 5 μm. c) Quantitation of FRET/Tq ratio from TqLckV2.3 biosensor in OT-I T WT cell hybridoma treated with vehicle (blue, n= 62) 10 μM PP2 Lck inhibitor (purple, n=60) or 2 min stimulation with 1 mM Pervanadate (green, n=65). ***, p < 0.001 by 1-way ANOVA with Tukey’s post hoc test.

Previous work has demonstrated that second-generation Lck FRET biosensors, including the design on which TqLckV2.3 is based, exhibit subcellular localization patterns comparable to wild-type Lck and accurately report conformational states associated with active, inactive, primed, and double-phosphorylated species [11, 15]. These findings support the physiological relevance of the biosensor and its ability to reflect endogenous Lck behavior.

We first confirmed the biosensor’s localization by confocal microscopy, we observed the characteristic localization of Lck at the plasma membrane and in a small intracellular vesicular pool, which was also previously described [18, 19] (Fig. 1b). Upon treatment with the Lck inhibitor PP2, we observed an increase in the FRET/Tq ratio, consistent with a closed (inactive) Lck conformation. Conversely, pervanadate stimulation caused a decrease in the FRET/Tq ratio because, in addition to inhibiting phosphatases, recent evidence shows that pervanadate directly activates Src-family kinases, including Lck, independently of TCR engagement [20]. Thus, the observed FRET decrease (Fig. 1c) reflects a chemically induced opening of the Lck biosensor rather than a simple mimic of physiological TCR triggering. Notably, unstimulated cells exhibited a broad distribution of FRET/Tq ratios, likely reflecting the presence of distinct Lck pools in different conformational states.

### Quantitative Analysis of Lck Regulation via TCR–pMHC Conjugate Formation in antigen presenting surfaces

To investigate Lck regulation in the context of antigen presentation, we developed an improved assay system tailored to measure T-cell receptor (TCR) engagement with pMHC complexes. The OT-I TCR recognizes the OVA peptide (SIINFEKL) with high affinity when presented by H-2K^b^ but does not respond to the VSV peptide (RGYVYQGL) in the same context. H-2K^b^-VSV, on its own is non-stimulatory, but can recruit CD8 to the synapse [21]. Specifically, we analyzed conjugate formation between OVAαβ cells expressing CD8α, CD8β and TqLckV2.3 (hereafter referred to as OT-I WT) and a surrogate antigen-presenting cell line. We used CHO cells engineered to express DOX inducible single-chain H-2K^b^ molecules constitutively loaded with either the agonist peptide OVA (scH-2K^b^–OVA) or the non-stimulatory peptide VSV (scH-2K^b^–VSV), as previously described [21, 22] (Fig. 2a, b). To generate stable and functional single-chain pMHCI (sc-pMHCI) molecules, we employed recombinant constructs in which the MHC-binding peptide was fused to the N-terminus of β2-microglobulin, followed by linkage to the N-terminus of the MHCI heavy chain, as previously described [23–25]. After DOX treatment, the constructs were stably expressed on the cell surface and were able to efficiently trigger T-cell activation (Fig. 2c).

**Figure 2.**
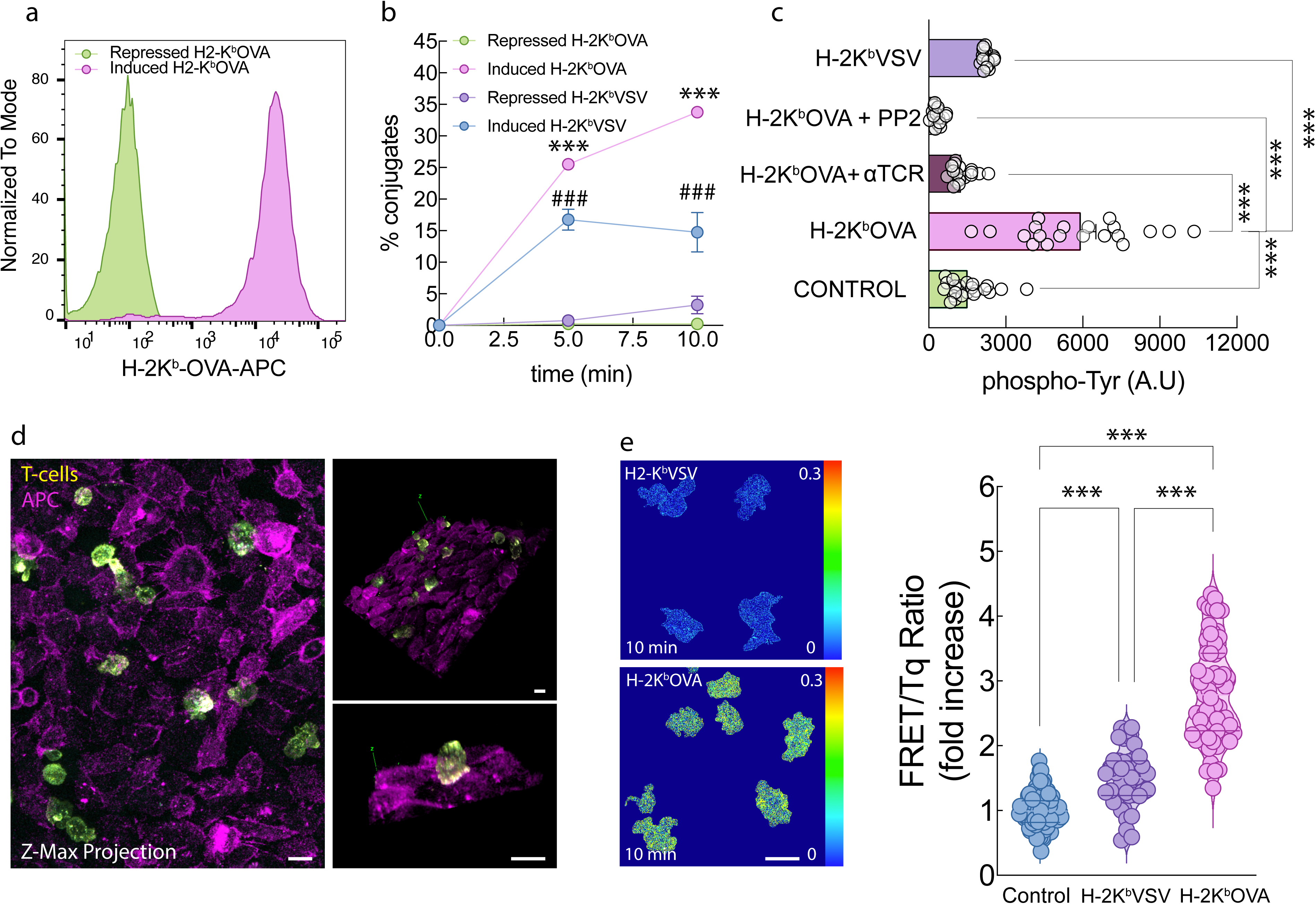
Expression of inducible sc-MHCI agonist molecules H-2K^b^-OVA in CHO cells and activation of corresponding OT-I CD8 T cells. a) Trex CHO cells expressing DOX-inducible sc-MHC were stained with anti-H-2K^b^-OVA and analyzed by flow cytometry. Green shading: Repressed (no DOX) Trex CHO cells. Purple: Induced (+ DOX) H-2K^b^-OVA. b) OT-I WT hybridomas expressing TqLckV 2.3 biosensor were stimulated for the indicated times with Cy5 labeled Trex CHO cells expressing DOX-inducible sc-MHC and percentage of conjugates were represented.). # Repressed H2-K^b^-VSV vs Induced H-2K^b^-VSV, p < 0.1; *** Repressed H2-K^b^-OVA vs Induced H-2K^b^-OVA, p < 0.001 by Student’s t test. c) OT-I WT hybridomas were stimulated as in b) for 15 min and stained with anti-phospho-Tyr. Images were taken using confocal microscopy. Mean fluorescence of SUM projections of Control n = 23, induced H-2K^b^-OVA n = 20, induced H-2K^b^-OVA + αTCR n= 18, induced H-2K^b^-OVA +PP2 n= 18 and induced H-2K^b^-VSV n = 17 ± s.e.m. is shown. ***, p < 0.001 by 1-way ANOVA with Tukey’s post hoc test. d) OT-I WT T cell hybridoma expressing TqLckV2.3 biosensor cells (yellow) were plated onto a Cy5 labeled monolayer of Trex CHO cells (purple) expressing DOX-inducible sc-MHC and analyzed by confocal microscopy. A representative Z-max projection image is presented, along with two distinct 3D reconstructions illustrating T cell–APC interactions. e) Individual FRET/Tq ratio (representative images shown) of OT-I WT T cell hybridoma expressing TqLckV2.3 cells. Control n = 98, induced H-2K^b^-OVA n = 65 and induced H-2K^b^-VSV n = 43. ***, p < 0.001 by 1-way ANOVA, multiple comparisons with Tukey’s post hoc test. Scale bar 5 μm.

Conjugate formation was determined by flow cytometry with previously Cy5-labeled APCs. Cy5, in its NHS-ester version reacts with primary amines, such as those on lysine residues or the N-terminus of proteins. It’s not cell-permeable what makes ideal for unspecific cell surface staining in live and fixed cells. Cells were fixed and conjugate formation was assessed by counting the percentage of cell couples positive for both mVenus and Cy5. Quantification revealed a time-dependent increase in the percentage of OT-I WT cells expressing TqLckV2.3 biosensor forming conjugates with CHO cells upon induction of pMHC expression. This increase was significantly greater when CHO cells presented the antigenic OVA peptide compared to the non-antigenic VSV peptide. The weak but distinct conjugate formation of H-2K^b^-VSV suggested that nonstimulatory (endogenous) pMHC complexes could be important for initial formation of T cell–APC conjugates. In the absence of DOX (repressed state), there was some “leaky” expression of H-2K^b^-OVA, which was sufficient to trigger partial CD69 upregulation in OT-I WT T cells (data not shown), however, this level of expression did not lead to significant TCR downregulation (12,13). Flow cytometry using a H-2K^b^-OVA-specific antibody revealed that H-2K^b^-OVA expression under these conditions was nearly undetectable (Fig. 2a).

To assess early T-cell activation, we measured global tyrosine phosphorylation levels after 15 minutes of co-culture using phospho-tyrosine staining (Fig. 2c). OT-I WT cells exposed to CHO cells presenting H-2K^b^-OVA exhibited elevated whole cell phospho-tyrosine levels relative to those exposed to H2-K^b^-VSV. Pre-treatment of OT-I WT cells with either a TCR-blocking antibody (Biolegend, clone B20.1) or the Src-family kinase inhibitor PP2 abrogated this phosphorylation response, reducing it to levels comparable to those observed with the non-stimulatory VSV peptide (Fig. 2c). Together, these results demonstrate that OT-I WT T cells selectively form stable conjugates and initiate early activation signaling in response to antigenic pMHC complexes, highlighting the specificity and functional sensitivity of TCR–pMHC interactions in this system.

### Validation of an improved second-generation FRET-based Lck biosensor for live-cell imaging

Having validated the TqLckV2.3 biosensor for monitoring Lck conformational changes, we next employed confocal microscopy to assess Lck dynamics in a more spatially resolved context. OT-I T cells expressing the biosensor were seeded onto monolayers of Cy5 labeled CHO cells inducibly expressing pMHC complexes. Z-projection imaging was used to capture the whole dimension between T cells and the CHO monolayer interaction as shown in Fig. 2d. Unexpectedly, when CHO cells presented the antigenic OVA peptide, we observed a significant increase in the FRET/Tq ratio across the entire T cell (Fig. 2e), suggesting a shift toward a closed (inactive) Lck conformation.

To explore this unexpected observation, we assessed Lck phosphorylation using phospho-specific antibodies by flow cytometry. Stimulation with anti-CD3/CD8 crosslinking induced a progressive increase in pY^505^-Lck, whereas total Lck remained constant (Fig. 3a). Quantification of the pY^505^/total Lck ratio confirmed a time-dependent rise, reaching ∼1.7-fold at 10 min post-stimulation (Fig.3b). Specificity of the phospho-Y^505^ staining was verified by PP2 treatment; PP2 elevated pY^505^–Lck signal intensity (Fig. S3), as anticipated for Lck inhibition. Conversely quantification of pY^394^-Lck/non-pY^394^-Lck showed no global increase but a slight decrease in pY^394^-Lck over time showing that pY^394^-Lck would remain membrane-enriched. When PP2 was present throughout the experiment (30-min pretreatment and during TCR stimulation), pY^394^ remained suppressed at all time points, consistent with complete inhibition of Lck catalytic activity. In contrast, pY^505^ increased over time despite PP2, reflecting unopposed Csk activity and CD45 exclusion at sites of TCR engagement. Thus, PP2 locks Lck in a persistently inactive, Y^505^-phosphorylated state even during potent TCR/CD8 stimulation (Fig 3a b).

**Figure 3.**
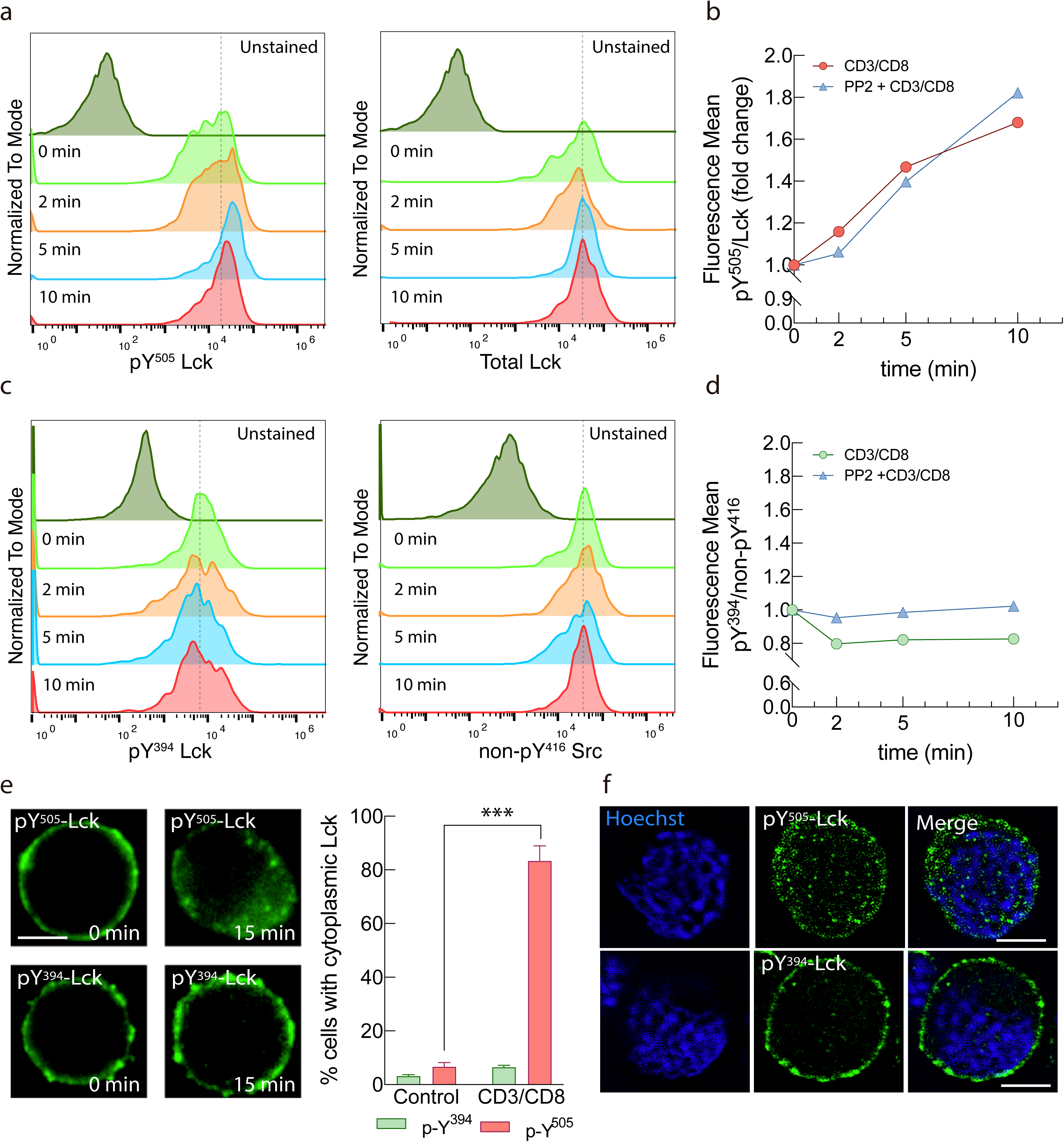
Spatial segregation of different endogenous phospho-Lck species. a) Flow cytometry histograms showing phosphorylation of Lck at Y^505^ (pY^505^, left) and total Lck (right) in OT-I WT T cells stimulated with anti-CD3/CD8 crosslinking for the indicated times. Secondary only controls (no primary antibody) are shown in gray. b) Quantification of pY^505^/total Lck mean fluorescence ratio over time in untreated (red) or pretreated with 10 μM PP2 (blue) for 20 min prior to stimulation with anti-CD3/CD8 crosslinking, expressed as fold change relative to t = 0 min. c) Flow cytometry histograms showing phosphorylation of Lck at Y^394^ (pY^394^, left) and non phospho Y^416^ Src (right) in OT-1 WT T cells stimulated with anti-CD3/CD8 crosslinking for the indicated times. Secondary only controls (no primary antibody) are shown in gray. d) Quantification of pY^394^/nonpY416 Src mean fluorescence ratio over time in untreated (green) or pretreated with 10 μM PP2 (blue) for 20 min prior to stimulation with anti-CD3/CD8 crosslinking, expressed as fold change relative to t = 0 min. e) OT-I WT T cells, unstimulated and 15 min after stimulation with anti-CD3/CD8 crosslinking was analyzed by epifluorescecnce deconvolution microscopy. Green: phospho-Lck antibody staining dsitribution analysis. Green: phospho-Lck antibody staining. >40 cells/sample. Nuclei were counterstained with Hoechst 33342. Bars represent % of cytoplasmic phospho-Lck ± s.e.m. f) Confocal representative section of OT-I WT T cells 15 min after stimulation with anti-CD3/CD8 crosslinking and analyzed by confocal deconvolution microscopy. ***, p < 0.001 by Student’s t test. Scale bar 5 μm.

Because staining was performed on fixed and permeabilized cells, we next examined whether this increase reflected enhanced phosphorylation or intracellular redistribution in stimulated cells. Immunofluorescence and confocal microscopy revealed that following TCR engagement, active Lck (pY^394^) remained at the plasma membrane, while inactive Lck (pY^505^) was predominantly internalized (Fig. 3e, d and f). These findings support the FRET-based observation of increased closed conformation at the whole-cell level and indicate that internalization of inactive Lck contributes to the overall FRET signal. To address whether TqLckV2.3 measurements reflects the same spatial partitioning we next tested the sensor under conditions where endogenous phospho-Lck species segregate. We directly compare the intracellular localization of the inactive Lck pool by phosphospecific staining in cells expressing TqLckV2.3 sensor and performed confocal imaging of the sensor with phospho-Y^505^ after stimulation. Line-scan profiles confirmed intracellular enrichment of pY^505^-Lck, reinforcing selective partitioning of inactive Lck. (Fig. S4)

The selective internalization of Y^505^-phosphorylated (inactive) Lck following TCR engagement suggests a previously unrecognized regulatory mechanism that may spatially restrict Lck activity by compartmentalizing its inactive pool away from the site of TCR stimulation, normally the immune synapse. Altogether, these findings establish TqLckV2.3 as a powerful biosensor for dissecting Lck conformational dynamics in live cells and uncover a novel mechanism of spatial regulation whereby inactive Lck is selectively internalized following TCR engagement, potentially fine-tuning signaling at the immune synapse.

### Lck biosensor reveals conformational activation at the immune synapse upon TCR engagement and distinguishes CD8 independent Lck pools

To investigate the spatial and conformational dynamics of Lck activation during TCR engagement we used fast FRET live cell microscopy. In order to test the system, stimulation with crosslinking biotinylated anti-CD3/CD8 antibodies was performed. We observed immediately after streptavidin addition a 5 minute decrease in the FRET/Tq ratio as shown in Fig. 4a. This conformational change was abolished by PP2 pre-treatment, it’s important to note that both traces include anti-CD3/CD8 stimulation, so the expected decrease in FRET is counterbalanced by the increase caused by PP2, keeping the signal relatively stable overall confirming that the observed FRET signal corresponds to Lck activation. We detected a pronounced difference in the global FRET response when comparing CD3/CD8 antibody crosslinking (Fig. 4a) to antigen presentation by CHO monolayers (Fig. 2e). Antibody-mediated crosslinking triggered a synchronous and widespread activation of Lck across the entire plasma membrane, reflected by an immediate decrease in the FRET/Tq ratio as the biosensor adopted an open conformation. This global activation is consistent with extensive Y^394^ phosphorylation throughout the cell surface [26, 27]. In contrast, stimulation with OVA-presenting CHO cells produced an asynchronous response, as individual T cells engaged the monolayer at different times and formed spatially restricted immune synapses. Under these physiological conditions, Lck activation was confined to the synapse, while the majority of Lck outside this region remained phosphorylated at Tyr^505^ and inactive. This pattern contributed to a net increase in whole-cell FRET measurements (Fig. 2e), consistent with previous reports that antigen-driven TCR triggering can promote inhibitory Y^505^ phosphorylation as part of feedback regulation of Lck activity [27]. To assess the biosensor’s performance in a more physiological context, we co-cultured TqLckV2.3-expressing OT-I WT cells with CHO cells expressing either H-2K^b^–OVA or H-2K^b^–VSV and monitored FRET changes at the immune synapse. A significant decrease in the FRET/Tq ratio was observed only in response to OVA presentation, indicating Lck activation in the immune synapse upon antigen recognition (Fig. 4b).

**Figure 4.**
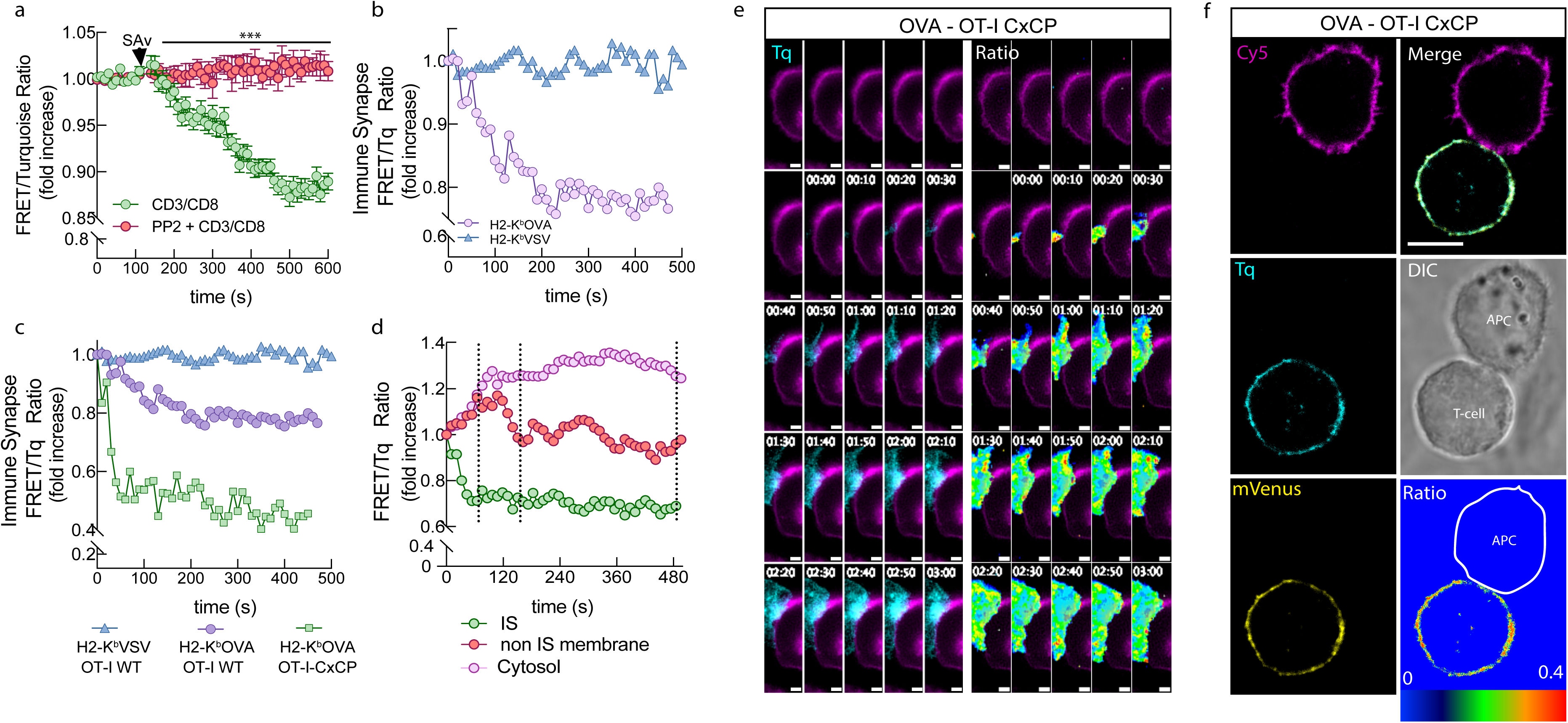
Lck dynamics at the Immune Synapse. a) OT-I WT T cells expressing TqLckV2.3, untreated (green) or pretreated with 10 μM PP2 (red) were stimulated both with biotinylated anti-CD3/CD8, 1 mg/ml streptavidin was added at the indicated time (arrow) and analyzed by live whole cell FRET microscopy. b) OT-I WT hybridomas expressing TqLckV2.3 were stimulated with Trex CHO cells expressing DOX-inducible H-2K^b^-OVA (purple circles) or H-2K^b^-VSV (blue triangles) and FRET/Tq ratio at the contact interface was represented since the initial contact between T cell and APC. c) Same as in B but with OT-I CxCP mutant (green squares). d) Differential FRET/Tq ratio analysis of OT-I CxCP cells trace from c) at the IS (green circles), the rest of the plasma membrane (red circles) and the cytosolic compartment (purple circles). The results are representative of at least three independent experiments. e) Detail of the contact zone of a time-lapse Tq fluorescence (left) or FRET/Tq ratio (right) images of OT-I CxCP T cells forming an immune synapse with Cy5 labeled CHO (purple) cells presenting OVA peptide. Time 0 of interaction is marked. f) Representative image of OT-I CxCP TqLckV2.3 expressing cell forming the immune synapse (Tq fluorescence in cyan, mVenus in yellow, CHO APC membrane labelled with Cy5 in purple and transmitted light image is shown as well) Bottom-right panel shows FRET/Tq ratio of the T cell. The results are representative of at least three independent experiments.

Previous results indicated that free Lck is initially responsible for TCR triggering [6]. To further dissect the contribution of CD8-bound versus free Lck, we expressed the TqLckV2.3 biosensor in OT-I.CxCP cells carrying a mutation in the intracellular CxCP motif of CD8α (C^227^K/C^229^P mutated to S^227^K/S^229^P), which disrupts Lck binding [6]. Interestingly, these mutant cells exhibited a more pronounced FRET/Tq ratio decrease upon OVA recognition compared to CD8 wild-type cells, suggesting that free Lck undergoes more substantial conformational changes than CD8-bound Lck during TCR engagement (Fig. 4c, e, f and Figs. S5 and S6). These results align with previous findings that initial TCR triggering is mediated predominantly by free Lck rather than CD8-bound Lck [6].To further investigate the spatial dynamics of free Lck during antigen presentation, we manually quantified FRET/Tq ratio of the Tq-LckV2.3 biosensor at the IS and across the remaining plasma membrane (Fig. 4d). During the initial phase of T cell–APC contact (0–60 s), we observed a marked decrease in FRET signal at the IS, indicative of sensor opening and most likely a higher Lck activity. In contrast, an increase in FRET was detected in the rest of the membrane, suggesting Lck inactivation in those regions. In the subsequent 60 seconds, the sensor at the rest of the membrane exhibited a closure, consistent with decreased Lck activity, which remained stable throughout the rest of the interaction. In parallel, the cytosol exhibited a sustained increase in the ratio that remained elevated throughout the recording. Notably, the kinetics of Lck conformational changes at the synapse and non-synapse were not perfectly synchronous. While Lck opened rapidly at the IS during the first ∼60 s, the non-synaptic membrane showed a slightly prolonged increase in FRET (closure) that extended for ∼120 s. This minor temporal discordance likely reflects the asymmetric spatial organization of early TCR signaling: rapid CD45 exclusion and local activation at the IS, combined with slower feedback processes such as Csk-mediated Y^505^ phosphorylation and internal redistribution of inactive Lck outside the synapse (see Fig. 3e, f and S4). After this short divergence, the non-synaptic regions returned to baseline, whereas Lck at the IS remained in an open conformation. These observations underscore the compartmentalized and temporally staggered nature of Lck regulation during immune synapse formation

### High-Resolution Imaging Reveals Spatial Restriction of Active Lck at the Immune Synapse

Since free Lck seems to be responsible for initial TCR triggering, to enhance spatial resolution and better characterize Lck conformational dynamics at the immune synapse (IS), we performed deconvolution confocal microscopy on fixed cells. Fig. 4f shows a representative conjugate between an OT-I CxCP mutant T cell expressing the TqLckV2.3 biosensor, where Lck cańt bind to CD8, and a Cy5-labeled CHO antigen-presenting cell expressing H-2K^b^-OVA (Fig. 4f).

FRET/Tq ratio imaging revealed a distinct decrease in the ratio at the IS upon recognition of H-2K^b^-OVA (Fig 4 e and f) (but not H-2K^b^-VSV; Fig 4b and c), indicating the presence of Lck in an open (active) conformation upon cognate antigen recognition. Notably, a subtle but consistent increase in the FRET/Tq ratio was observed at the periphery of the IS. This is consistent with the elevated FRET/Tq levels previously detected in the non-synaptic membrane regions by live-cell fluorescence microscopy (Fig. 4d). This spatial pattern suggests that only the open conformation of Lck is selectively enriched within the synapse, while the closed (inactive) form may be excluded or retained at the borders.

### Csk Preferentially Phosphorylates Lck not bound to CD8

To establish a latent system where we can distinguish the role of spatial conformation of free Lck and CD8 bound Lck, we next sought to restrain the kinase activity of Lck by overexpressing C-terminal Src kinase (Csk). Csk is a key negative regulator of Lck, phosphorylating its inhibitory tyrosine residue (Y^505^) to promote a closed, inactive conformation [28–30]. This well-known phosphorylation event counterbalances the activating phosphorylation at Y^394^, thereby modulating Lck catalytic activity. Importantly, Csk lacks intrinsic membrane-targeting domains and relies on adaptor proteins such as PAG/Cbp to localize to the plasma membrane, where it can access Lck. To investigate whether Csk preferentially phosphorylates CD8-bound or free Lck, we expressed the FRET-based Lck activity sensor Tq-LckV2.3, along with Csk and Cbp, which increases the availability of raft-localized docking sites for Csk [31], in OT-I WT or OT-I CxCP cells where Lck–CD8 interaction is broken. In cells expressing the CxCP mutant, we observed a marked increase in the FRET/Tq ratio of the Lck sensor when Csk and Cbp were co-expressed, indicative of a closed Lck conformation and consistent with enhanced phosphorylation at the inhibitory Y^505^ site (Fig. 5d). These results suggest that Csk preferentially targets free Lck for inhibitory phosphorylation. Interestingly, Csk was comparably overexpressed across both cell lines (Fig. 5a), ruling out expression imbalance as the basis for the differential FRET response.

**Figure 5.**
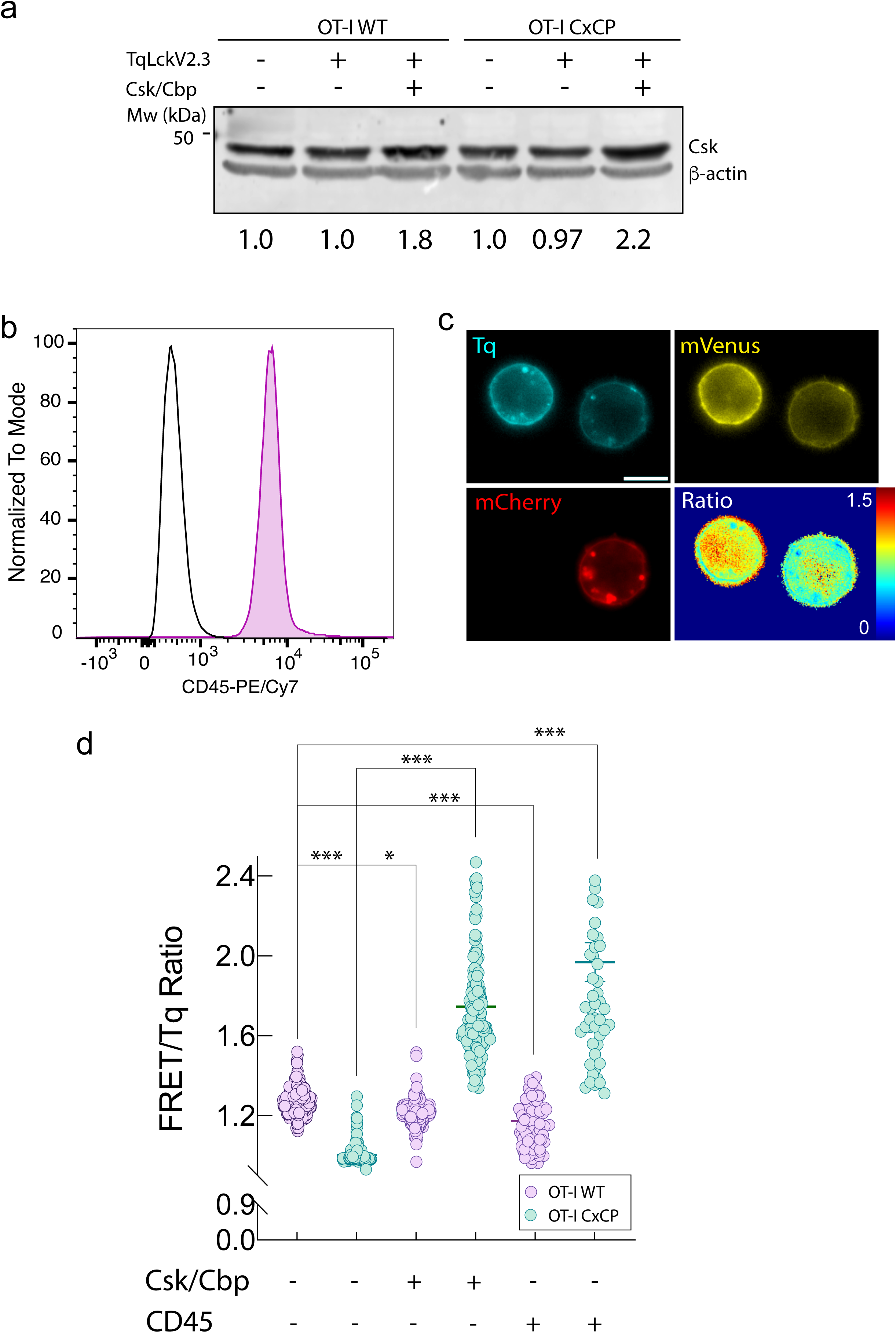
Lck regulation in resting cells. a) Csk overexpression. OT-I WT and OT-I CxCP cells expressing TqLckV2.3 sensor and transduced or not with Csk-P2A-Cbp as indicated were analyzed for Csk expression by immunoblot. β-actin was used as loading control. Signal intensity ratios of Csk/β-actin were calculated and is shown beneath b) Endogenous CD45 expression. Flow cytometry histogram of CD45-PE/Cy7 staining in OT-I WT T-cells (purple) compared with isotype control (black). Histograms are normalized to mode. c) Representative imaging of CD45 overexpression. Confocal image of OT-I WT cells expressing TqLckV2.3 and either transduced with CD45cytCD43ex-mCherry (red) or untransduced (no red). FRET/Tq ratio is shown as a pseudocolor map for each cell. Scale bar, 10 μm. d) Quantification of Lck conformation. FRET/Tq ratio from TqLckV2.3 biosensor in unstimulated OT-I WT (purple) or OT-I CxCP (cyan) cell hybridomas expressing Csk/Cbp or CD45_cyt_CD43_ex_. WT n=233, CxCP n=139; Csk-Cbp expressing WT n=262 CxCP Csk-Cbp CxCP n=149; CD45 expressing WT n=119 CD45 expressing CxCP n =52. *, p < 0.05; ***, p < 0.001 by 1-way ANOVA, multiple comparisons with Tukey’s post hoc test.

### CD45 Suppresses Free Lck Activation via Y^394^ Dephosphorylation to Prevent Spontaneous TCR Signaling

CD45 is a tyrosine phosphatase that modulates T cell signaling in a complex manner by dephosphorylating both the inhibitory Y^505^ and the activating Y^394^ of Lck, thereby exerting both activating and suppressive control depending on the context and localization of Lck. The balance between these opposing activities shifts according to the amount of CD45 available. At low levels, CD45 primarily removes pY^505^ and promotes Lck activation, whereas at higher levels CD45 increasingly targets the activating pY^394^ residue, resulting in net inhibition of Lck activity. To investigate the regulatory role of CD45 on distinct pools of Lck in resting T cells, we examined the conformational changes in free (non CD8-bound) Lck. We made use of the construct CD45cytCD43ex-mCherry [32, 33] where the cytoplasmic phosphatase domains of CD45 were fused to the large extracellular and transmembrane domains of CD43. Although CD43 ectodomain shares no amino acid similarity with CD45, it is as long and as heavily glycosylated as that of CD45 and should therefore mimic its basic physicochemical properties. This construct has previously been shown to functionally replicate CD45 [32, 33]. Thus, the CD45cytCD43ex construct was used not to restore CD45 function, but to increase CD45 phosphatase activity above endogenous levels in a controlled manner. Flow-cytometric staining confirmed that OT-I cells endogenously express CD45 at high levels (Fig. 5b), as expected for T-lineage cells, and therefore CD45cytCD43ex allows us to test how elevated CD45 phosphatase availability modulates Lck regulation beyond endogenous levels.

We used the TqLckV-2.3 biosensor in the presence or absence of CD45_cyt_CD43_ex_-mCherry in resting cells, in OT-I WT cells, CD45cytCD43ex had a modest, but significant, tendency to lower FRET/Tq ratio, consistent with partial removal of inhibitory pY^505^ on CD8-bound Lck (Fig. 5c, d). In contrast, we observed a consistent FRET increase in the free Lck pool when CD45_cyt_CD43_ex_-mCherry was expressed in OT-I CxCP cells (Fig. 5d). This suggests that CD45 preferentially targets Y^394^ in free Lck, in contrast to its well-characterized role in dephosphorylating the inhibitory Y^505^ site as shown when Lck can be also bound to CD8 (Fig. 5c, d). These findings imply that CD45 may act as a negative regulator of free Lck by maintaining it in an inactive state through dephosphorylation of the activating site. This observation suggests that CD45 actively suppresses the activation of free Lck by targeting its activating phosphorylation site. It is well stablished that CD45 functions as a gatekeeper, but the results suggest that particularly maintain free Lck in an inactive state under resting conditions. This mechanism may serve to prevent spontaneous or ligand-independent TCR triggering, ensuring that T cell activation is tightly regulated and spatially restricted to appropriate signaling contexts. The concept of CD45 as a rheostat for Lck activity is not novel to the present study; rather, it was very elegantly formulated by D. Alexander’s group [34]. The novelty on our research is that CD45 seems to have preference for free Lck instead CD8 bound.

## DISCUSSION

T cell activation is a tightly regulated process that relies on the precise spatial and temporal coordination of signaling molecules at the immune synapse. Among these, the Src-family kinase Lck plays a pivotal role in initiating TCR signaling by phosphorylating ITAMs within the CD3 complex. While the importance of Lck is well established, the mechanisms that regulate its activation and localization during antigen recognition remain incompletely understood. In this study, we employed an improved second-generation FRET-based biosensor, TqLckV2.3, to visualize Lck conformational dynamics in live T cells and dissect the contributions of CD8-bound versus free Lck during TCR engagement.

TqLckV2.3 biosensor reports on the conformational state of Lck but does not differentiate between the primed (open, non-phosphorylated) and double-phosphorylated (open, phosphorylated at both Y^394^ and Y^505^) forms, as both adopt a constitutively open conformation that is insensitive to TCR stimulation [8, 35]. The quantification of constitutively active Lck in resting T cells has been marked by striking controversy, with reported values ranging from ∼2% to ∼40% of the total Lck pool—a 20-fold discrepancy that has significant implications for our understanding of TCR signaling initiation [8, 9]. This dramatic variation appears to stem largely from methodological artifacts, particularly the spontaneous autophosphorylation of Lck that occurs after cell solubilization, as critically demonstrated by Ballek and colleagues who attributed much of the discrepancy to post-lysis phosphorylation events rather than true physiological activity levels [9]. The importance of controlling for this artifact has been further validated in Wei et al., where we addressed the issue by including the SFK inhibitor PP2 during cell lysis and immunoprecipitation procedures, thereby preventing artificial inflation of measured Lck activity [36]. Given that even carefully processed samples show functionally significant constitutive Lck activity—as evidenced by the correlation between Lck phosphorylation status and cytotoxic function in memory T cell subsets [10] future studies investigating steady-state Lck activity should, at minimum, include kinase inhibitors such as PP2 during sample processing to ensure accurate quantification and avoid the methodological pitfalls that have confounded this field.

Recent advances in fluorescence-based imaging techniques, particularly the development of FRET-based biosensors, have opened new possibilities for investigating the spatial and temporal aspects of Lck activation [11, 15, 16]. While FRET-based methods avoid post-lysis artifacts that plague biochemical approaches, they require careful interpretation and validation with complementary techniques. These approaches allow for real-time monitoring of Lck conformational changes but not a direct measure of the activation states within the complex environment of the immune synapse, providing unprecedented insight into the spatial regulation of TCR signaling machinery.

As mentioned earlier, the TqLckV2.3 biosensor reports conformational changes rather than catalytic activity. A decrease in the FRET/Tq ratio indicates a transition to an open conformation, but this state can correspond to multiple biochemical forms of Lck, including active Lck phosphorylated at Y^394^, primed Lck that is open but not phosphorylated, and even double-phosphorylated Lck (Y^394^ and Y^505^). Consequently, while our data demonstrate spatial and temporal shifts in Lck conformation during CD8^+^ T cell activation, they do not allow us to directly quantify enzymatic activity or distinguish between these open states. This limitation should be considered when interpreting the functional significance of FRET changes, and future studies combining FRET imaging with phospho-specific probes or activity reporters will be necessary to resolve these distinctions. Our results demonstrate that Lck undergoes distinct conformational changes upon TCR stimulation, with a rapid decrease in the FRET/Tq ratio indicating a transition to an open conformation. This activation was abrogated by the Src-family kinase inhibitor PP2, confirming the specificity of the biosensor for Lck activity. Importantly, we observed that antigen-specific stimulation with CHO cells presenting OVA-loaded pMHC induced a robust FRET response, whereas non-stimulatory VSV-loaded pMHC did not, validating the biosensor’s sensitivity to physiologically relevant TCR signals.

Unexpectedly, whole-cell imaging revealed an increase in the FRET/Tq ratio upon antigen recognition, suggesting a paradoxical shift toward a closed Lck conformation. FACS and immunofluorescence analysis resolved this discrepancy by showing that inactive Lck (pY^505^) is selectively internalized following TCR engagement, while active Lck (pY^394^) remains at the plasma membrane. This spatial segregation implies a novel regulatory mechanism in which the internalization of inactive Lck may serve to sharpen the signaling interface by restricting kinase activity to the immune synapse.

While the intracellular enrichment of pY^505^-Lck is compatible with partial internalization of this inactive conformation, other mechanisms may act in parallel to generate the spatially segregated pools of Lck that we observe. Several studies have demonstrated that Lck trafficking is strongly influenced by lipid-binding chaperones that regulate the solubility and targeted delivery of the kinase. In particular, Unc119 serves as a selective solubilizing factor for myristoylated Lck and promotes the directed transport of active Lck toward the immunological synapse [37]. Earlier work similarly showed that Unc119 regulates Src-family kinase dynamics by extracting Lck from membranes and enabling its polarized redistribution during T cell activation [38]. These findings raise the possibility that differences in chaperone-mediated extraction, cytosolic mobility, or targeted delivery could contribute to the differential localization of pY^394^- and pY^505^-Lck. Thus, internalization of inactive Lck likely represents one component of a broader set of trafficking processes—including Unc119-dependent solubilization—that together shape the spatial partitioning of Lck conformations in T cells.

In addition, our use of a CD8α mutant that disrupts Lck binding revealed that free Lck undergoes more substantial conformational opening than CD8-bound Lck. This finding supports previous models suggesting that free Lck is primarily responsible for initiating TCR signaling, while CD8-bound Lck may play a more modulatory role [6, 36]. High-resolution imaging confirmed that opened Lck is enriched within the immune synapse, whereas the inactive form is excluded to the periphery, reinforcing the idea that spatial compartmentalization of Lck activity is critical for signal fidelity.

In mice, the presence of Lck bound to CD8 is not essential for the antiviral and antitumor functions of cytotoxic T cells. However, it plays a kinase-dependent role in enhancing CD8^+^ T cell responses to weaker antigens. Conversely, Lck associated with CD4 is crucial for the proper development and activity of helper T cells, primarily through a kinase-independent mechanism that stabilizes CD4 on the cell surface [4]. As free Lck is the primary initiator of TCR signaling in CD8 T cells [6], then the Csk preference for phosphorylating free Lck suggests a built-in regulatory checkpoint. This ensures that only the right spatial and temporal context allows TCR triggering, preventing spontaneous or inappropriate activation. The system may keep free Lck in a controlled but restrained state (most likely in a closed configuration), where it can rapidly respond to TCR engagement but is otherwise kept inactive by Csk. This would allow rapid signal initiation while maintaining immune tolerance. CD8-bound Lck might serve more as a stabilized, less dynamic pool, possibly involved in signal amplification rather than initiation. This division of labor could explain why CD8-bound Lck is less accessible to Csk. During antigen recognition at the IS, Lck remains in a sustained open conformation, supporting continuous TCR signaling. This localized activation ensures that signaling is tightly focused at the site of contact between the T cell and the antigen-presenting cell. In contrast, Lck outside the IS undergoes only transient inactivation, suggesting a dynamic but restrained regulation across the rest of the plasma membrane. This spatial compartmentalization of Lck activity likely serves to prevent unintended signaling while maintaining the cell’s readiness to respond to new stimuli, highlighting a finely tuned mechanism of T cell activation (Fig. 6).

**Figure 6.**
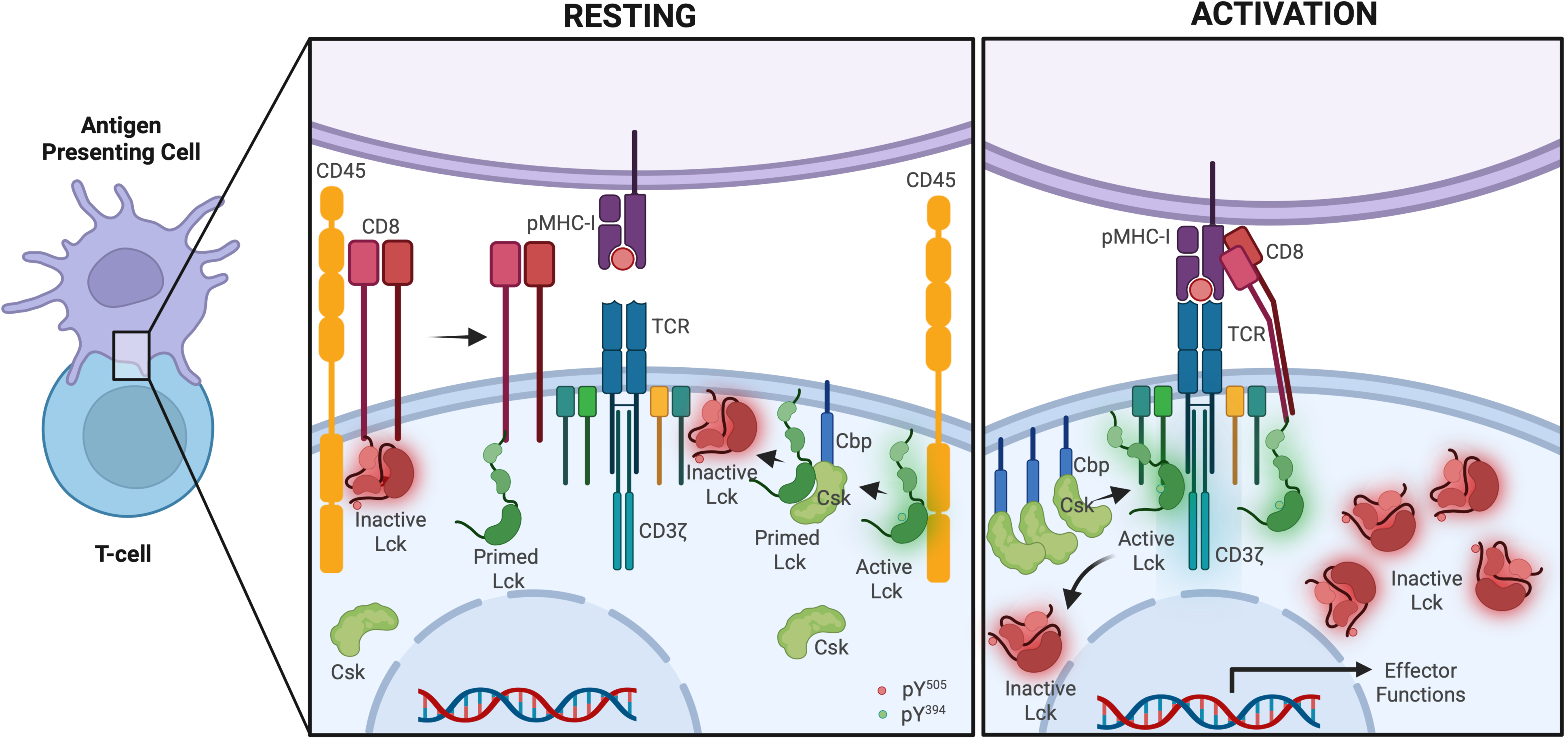
Spatial regulation of Lck and CD8 co-receptor dynamics in resting and activated CD8⁺ T cells. Schematic representation of the molecular organization and early signaling events at the CD8⁺ T-cell–APC interface under resting (left) and activating (right) conditions. **Resting state:** CD8, CD45, and the TCR–CD3 complex remain widely distributed across the membrane, with minimal physical proximity to pMHC-I on the antigen-presenting cell. Lck exists predominantly in inactive or primed conformations, controlled by opposing phosphorylation at Y^505^ (inhibitory) and Y^394^ (activating). Active regulation is mediated by Csk, recruited through Cbp/PAG microdomains, which phosphorylates Y^505^ to maintain Lck in its closed conformation. CD45 functions as a rheostat, continuously dephosphorylating Lck, but with substrate preference that depends on the context: at baseline, high CD45 accessibility maintains CD8-bound Lck in a primed, signaling-competent form by removing inhibitory pY^505^, while simultaneously suppressing spontaneous activation of free Lck by dephosphorylating pY^394^. This balanced, steady-state activity ensures low signaling noise while preserving readiness for antigen encounter. **Activation state:** Recognition of cognate pMHC-I induces co-aggregation of TCR–CD3 and CD8, facilitating recruitment of CD8-associated Lck and enabling rapid phosphorylation of CD3 ITAMs. The initial phosphorylation of the TCR is mediated by free Lck, which is the first pool able to access and phosphorylate CD3ζ upon antigen encounter before CD8-bound Lck is engaged and stabilizes TCR-pMHC-CD8 complex. Formation of the close-contact zone triggers size-dependent segregation of CD45, which is progressively excluded from the immune synapse due to its bulky extracellular domain. CD45 exclusion reduces its access to ITAMs and active Lck, permitting accumulation of phosphorylated CD3ζ and stabilization of active Lck at the synapse. Concurrently, Csk activity sustains phosphorylation of Y^505^, reinforcing spatial compartmentalization in which Lck adopts an open conformation exclusively at the immune synapse, while inactive Lck undergoes internalization. This orchestrated segregation of CD45, Csk, and distinct Lck pools sharpens the signaling interface and enables sustained antigen-driven effector responses. Created in https://BioRender.com.

It is important to highlight CD28-Lck interaction. CD28 doesn’t just “boost” signaling; it actively reorganizes signaling microclusters by tethering Lck. CD28 co-stimulation significantly increases the fraction of active Lck (pY^394^-Lck) localized in TCR microclusters compared to stimulation through TCR alone. This enhancement is not just recruitment but also involves greater spatial segregation between active and inactive Lck species, which lowers the activation threshold for downstream signaling (e.g., ZAP70 phosphorylation [39]. Although this is interesting in itself, the context of activation in that work is not the most physiological situation, as these observations were made by adding cells to surfaces coated with immobilized antibodies rather than using antigen-presenting cells, which provide a more natural signaling environment. Future studies could explore the impact of CD28 co-stimulation by engineering CHO antigen-presenting cells to express mouse CD28 ligands (e.g., CD80/CD86), thereby enabling a more physiologically relevant analysis of Lck dynamics under conditions that mimic full T cell activation.

Our data refine the current understanding of Lck regulation in resting T cells by demonstrating that Csk preferentially phosphorylates free, non-CD8-bound Lck, while CD45 dephosphorylates not only the inhibitory Y^505^ but also the activating Y^394^ residue. These findings challenge the notion of a static, constitutively active Lck pool and suggest a more dynamic regulatory landscape in which Lck activity is tightly controlled through spatial segregation and substrate specificity. In this context, CD45 emerges as a dual-function phosphatase capable of both priming and attenuating Lck activity depending on its localization and substrate accessibility. CD45 is a critical regulator that maintains T cells in a quiescent state and sets the threshold for TCR activation [40]. This nuanced model aligns with and extends the kinetic segregation framework, emphasizing the importance of membrane microdomain organization and the differential regulation of CD8-associated versus free Lck in shaping the threshold and timing of TCR signaling.

Together, these findings provide new insights into the spatial and conformational regulation of Lck during T cell activation. The selective internalization of inactive Lck and the preferential activation of free Lck at the CD8 immune synapse represent previously unrecognized layers of regulation that may fine-tune TCR signaling. The TqLckV2.3 biosensor offers a powerful tool for further dissecting these dynamics in physiological and pathological contexts, including autoimmunity and immunotherapy. Although our study focused exclusively on CD8⁺ T cells, we did not examine Lck dynamics in CD4⁺ T cells or assess how CD28 co-stimulation influences Lck regulation. These aspects should be explored in future studies. While the TqLckV2.3 biosensor could be applied to these contexts, technical constraints in our system prevented such analyses. Future work should address whether similar spatial and conformational regulation occurs in CD4^+^ T cells and during co-stimulatory signaling.

## DECLARATIONS

### Consent for publication

Not applicable.

### Availability of data and material

Available upon request.

### Author Contributions

Conceptualization, J.C. and C.M.; methodology, J.C. and C.M;.; validation, J.C. and C.M.; formal analysis, J.C. and G.S.J.; investigation, J.C.; resources, J.C and M.A.B.; data curation, J.C and G.S.J.; writing—original draft preparation, J.C.; writing—review and editing, J.C.; visualization, J.C.; supervision, J.C.; project administration J.C.; funding acquisition, J.C. All authors have read and agreed to the published version of the manuscript.

## Acknowledgments

We thank Montse Duque, Eva Merino and Álvaro Martín for excellent technical assistance. J.C thank Lorena, Manuela & Simón for their critical reading of the manuscript and specifically Nicholas Gascoigne for all the materials and advice provided through the year.

---

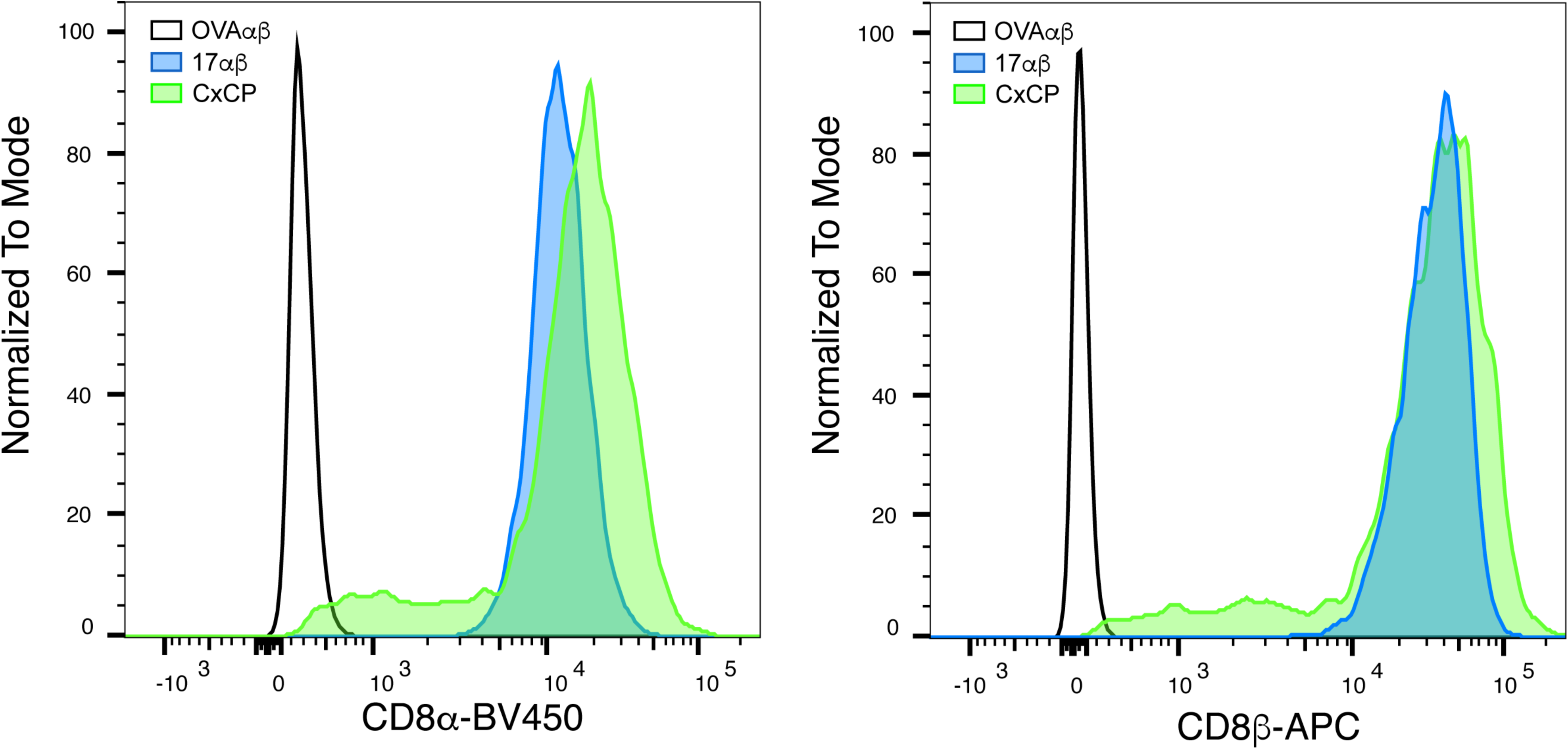

**FIGURE S2.**
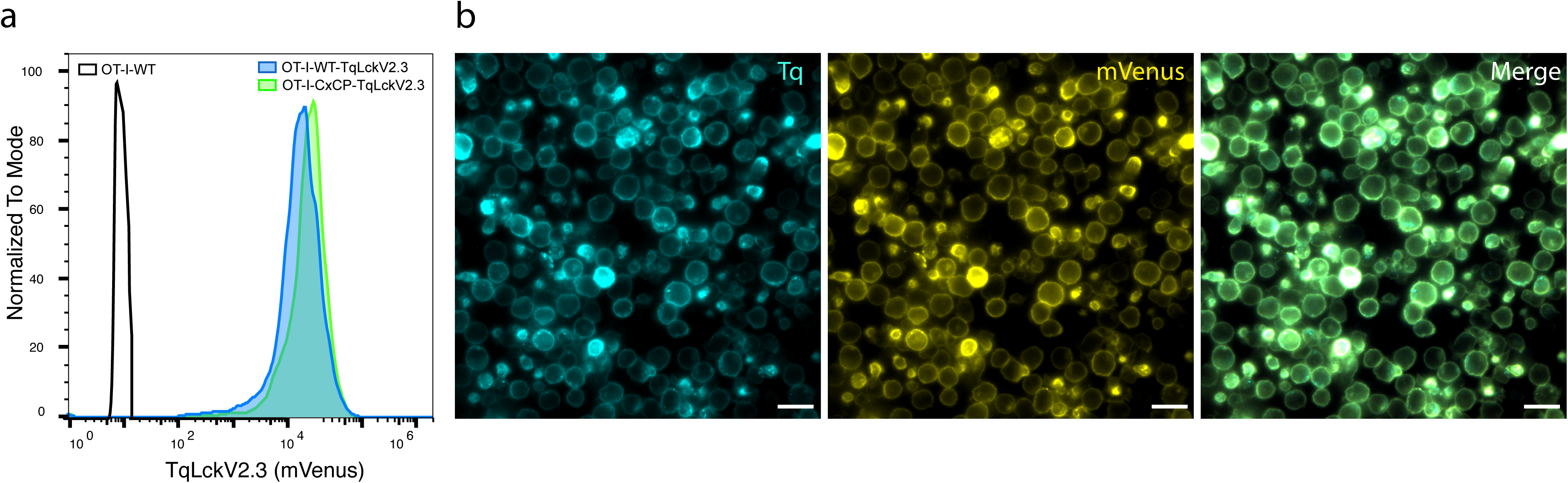

**FIGURE S3.**
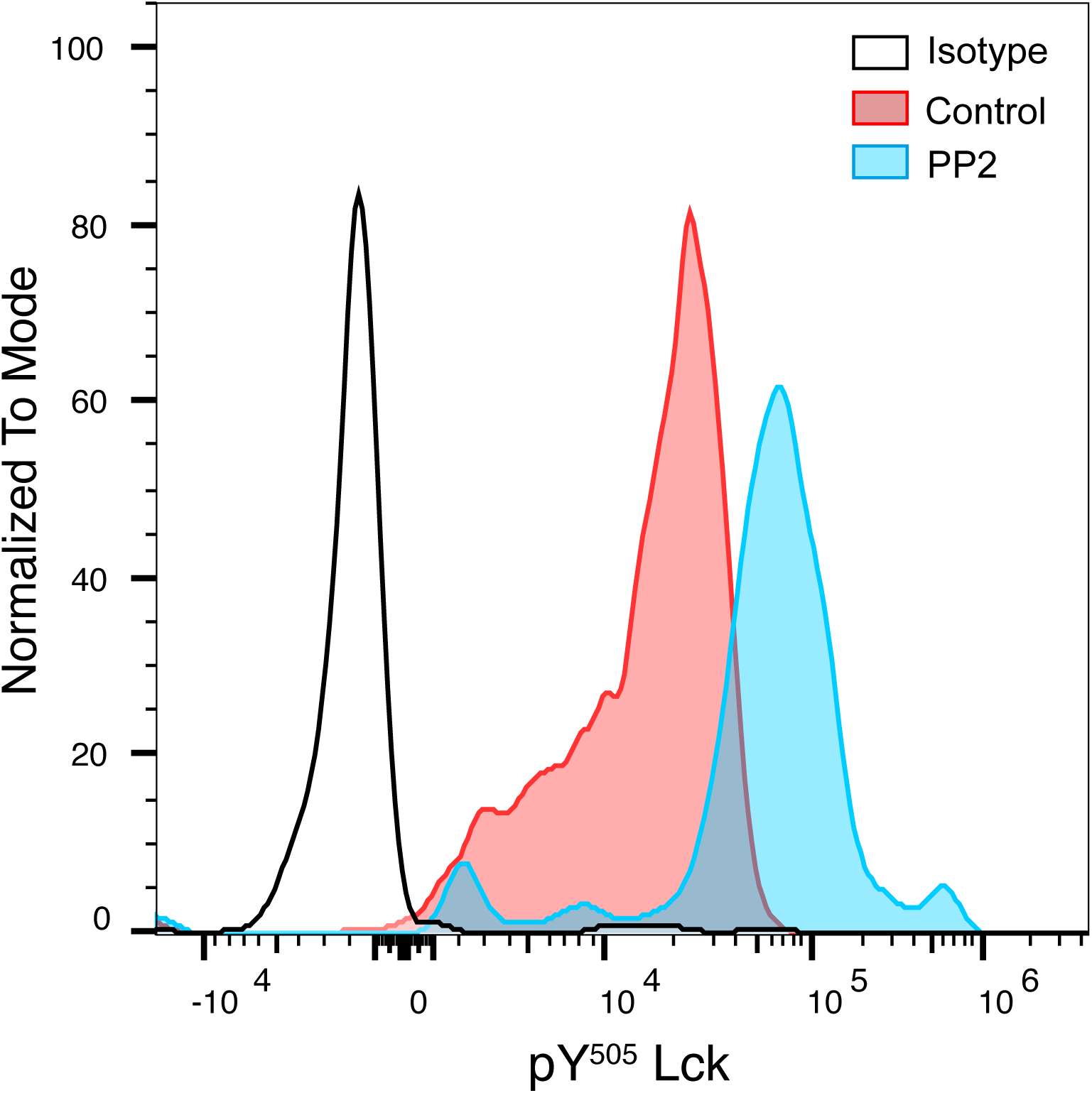

**FIGURE S4.**
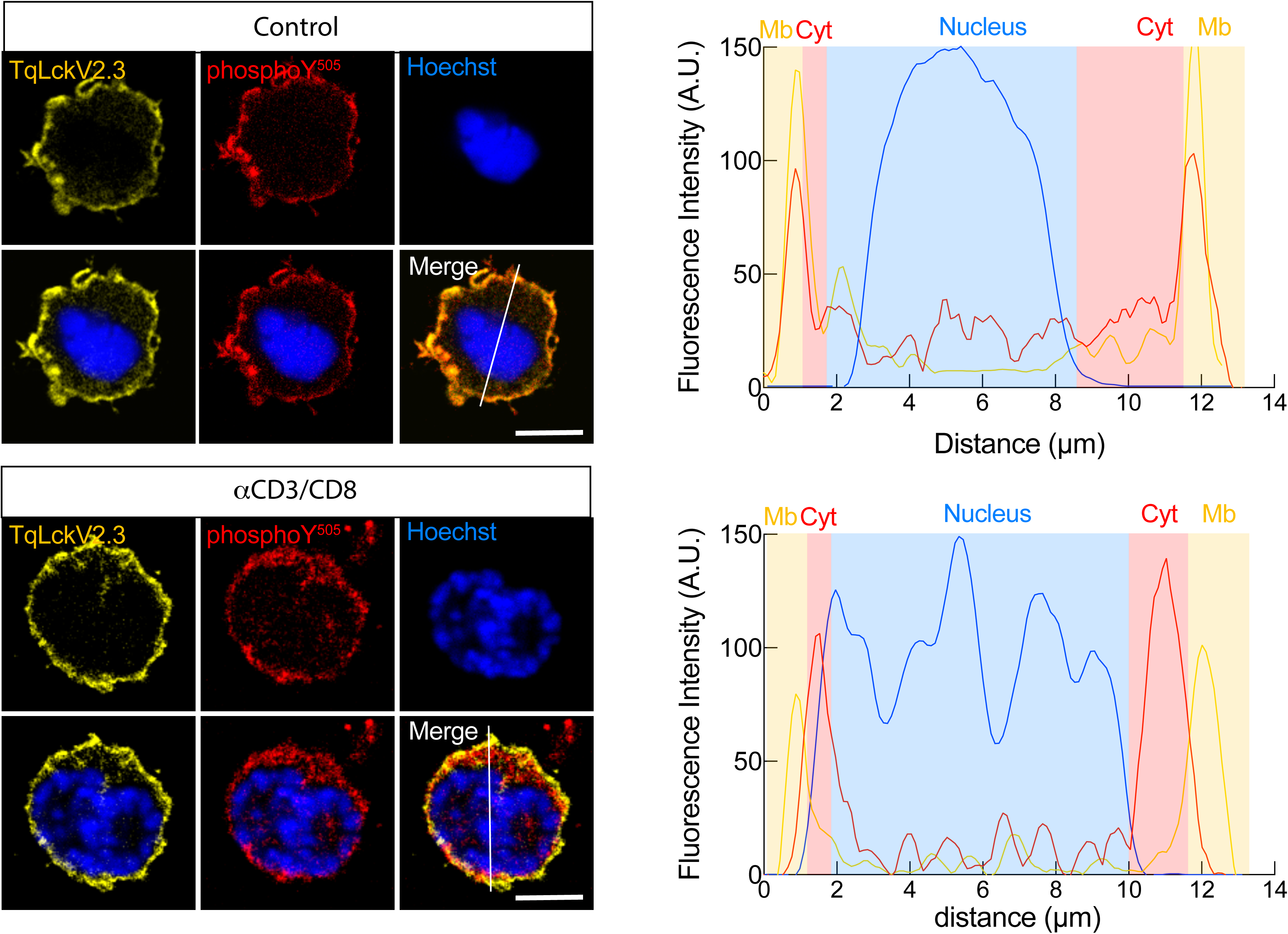

