## Supplementary Info for "Spatial Regulation of Lck Activation at the CD8 Immune Synapse Revealed by a FRET-Based Biosensor"

This PDF file includes:

Key Resources Table

Figure S1

Figure S2

Figure S3

Figure S4

Figure S5 (Movie)

Figure S6 (Movie)

### *Key Resources Table*

| Primary Antibodies | Source | Identifier/RRID |
|---|---|---|
| BV450 anti-CD8α (clone 53-6.7) | BD Biosciences | #560469/AB_1645281 |
| APC anti-CD8β (clone H35-17.2) | eBioscience | #17-0083-81/AB_657760 |
| PE/Cy7 anti-CD45 (clone 30F11) | BioLegend | # 103113 /AB_312978 |
| OVA257-264 (SIINFEKL) peptide bound to H-2Kb mAb (clone 25-D1.16) APC conjugated | Thermo Fisher Scientific | #175743-80/AB_1311288 |
| Purified anti-mouse TCR Vα2 | BioLegend | #127802/AB_1089248 |
| Biotin anti-mouse CD3ε (clone 145-2C11) | BioLegend | #100303/AB_312668 |
| Biotin anti-mouse CD8α (clone 53-6.7) | BioLegend | #100703/AB_312742 |
| Purified Mouse Anti-Csk (clone 52/Csk) | BD Biosciences | #610080/AB_397488 |
| Mouse Monoclonal Anti-β-Actin (clone AC15) | Sigma-Aldrich | #A5441/AB_476744 |
| Rabbit mAb Anti-Lck (clone 73A5) | Cell Signaling Technologies | #2787/AB_659970 |
| Rabbit anti-Lck Phospho-Y505 | Cell Signaling Technologies | #2751/AB_330446 |
| Rabbit mAb anti-Lck phospho-Y394 (clone E5L3D) | Cell Signaling Technologies | #70926/AB_2924371 |
| Mouse anti-Src non-phosphoY416 (clone 7G9) | Cell Signaling Technologies | #2102/AB_331358 |
| Mouse mAb Anti-phospho-Tyrosine (clone 4G10) | Millipore | #05-321/AB_309678 |
| **Secondary Antibodies** | **Source** | **Identifier/RRID** |
| F(ab')2-Goat anti-Rabbit IgG (H+L) Cross-Adsorbed Secondary Antibody, Alexa Fluor 488 | Thermo Fisher Scientific | #A11070/AB_2534114 |
| F(ab')2-Goat anti-Rabbit IgG (H+L) Cross-Adsorbed Secondary Antibody, Alexa Fluor 594 | Thermo Fisher Scientific | #A11072/AB_10563744 |
| F(ab')2-Goat anti-Mouse IgG (H+L) Cross-Adsorbed Secondary Antibody, Alexa Fluor 568 | Thermo Fisher Scientific | #A11019/AB_10563405 |
| Goat anti-mouse IgG IRDye 800CW | Li-Cor | #925-32210/AB_2687825 |

*SUPLEMMENTARY FIGURES*

**Figure S1. Surface expression of CD8α and CD8β on different T cell hybridomas.** Flow cytometry histograms showing normalized to mode fluorescence intensity for CD8α (left, BV450) and CD8β (right, APC) staining on three T cell hybridomas: OVAαβ (black), 17αβ (blue), and CxCP (green). Data indicate that OVAαβ cells exhibit negligible CD8 expression, whereas 17αβ and CxCP hybridomas display comparable and robust levels of both CD8α and CD8β chains.

**Figure S2. Expression of the TqLck-V2.3 FRET biosensor in OT-I hybridomas.** a) Flow-cytometry histograms of mVenus (biosensor acceptor) fluorescence for OT-I WT–TqLck-V2.3 (blue) and OT-I CxCP–TqLck-V2.3 (green), indicating matched expression levels across the two lines. The parental OT-I WT (no sensor) population (black, open) is shown as a negative control. Histograms are normalized to mode; cells used in all experiments were FACS-sorted through a narrow mVenus positive gate to ensure a uniform expression window prior to imaging and functional assays. b) Representative widefield images from the sorted populations acquired with identical settings illustrate homogeneous single-cell expression of Turquoise (cyan) and mVenus (yellow); the merge confirms co-expression across the field. Scale bars, 10 μm.

**Figure S3. Specificity of phsophoY$^{505}$ Lck staining.** Flow cytometry histograms showing normalized to mode fluorescence intensity for phosphoY$^{505}$ Lck staining on OT-I WT hybridoma before (red) and after (blue) 20 min treatment with 10 μM of PP2.

**Figure S4. Intracellular enrichment of pY505–Lck of the Lck FRET biosensor after TCR stimulation.** Representative confocal images of OT-I T-cell hybridomas expressing TqLckV2.3 (yellow), co-stained for phospho-Y$^{505}$ Lck (red) and Hoechst (blue). Cells were either unstimulated (Control) a) or stimulated by anti-CD3/anti-CD8 crosslinking (αCD3/CD8; 15 min) b). Right panels show line-scan profiles taken along the indicated white line, plotting fluorescence intensity for pY$^{505}$ (red), TqLckV2.3 (yellow), and Hoechst (blue) versus distance. Scale bar, 5 μm. Data are representative of 2 independent experiments.

**Figure S5. Detail of the immune synapse formation in OT-I CxCP T cell.** Time-lapse of Tq fluorescence (left) or FRET/Tq ratio (right) images of OT-I CxCP T cell forming an immune synapse with Cy5 labeled CHO (purple) cell presenting OVA peptide. Time 0 of interaction is marked.

**Figure S6. Immune synapse formation in OT-I CxCP T cell.** Time-lapse of Tq fluorescence (left) or FRET/Tq ratio (right) images of OT-I CxCP T cell forming an immune synapse with Cy5 labeled CHO (purple) cell presenting OVA peptide. Time 0 of interaction is marked.
