## Supplementary figures and images for "Spatial Regulation of Lck Activation at the CD8 Immune Synapse Revealed by a FRET-Based Biosensor"

### Supplemental Fig 1-4

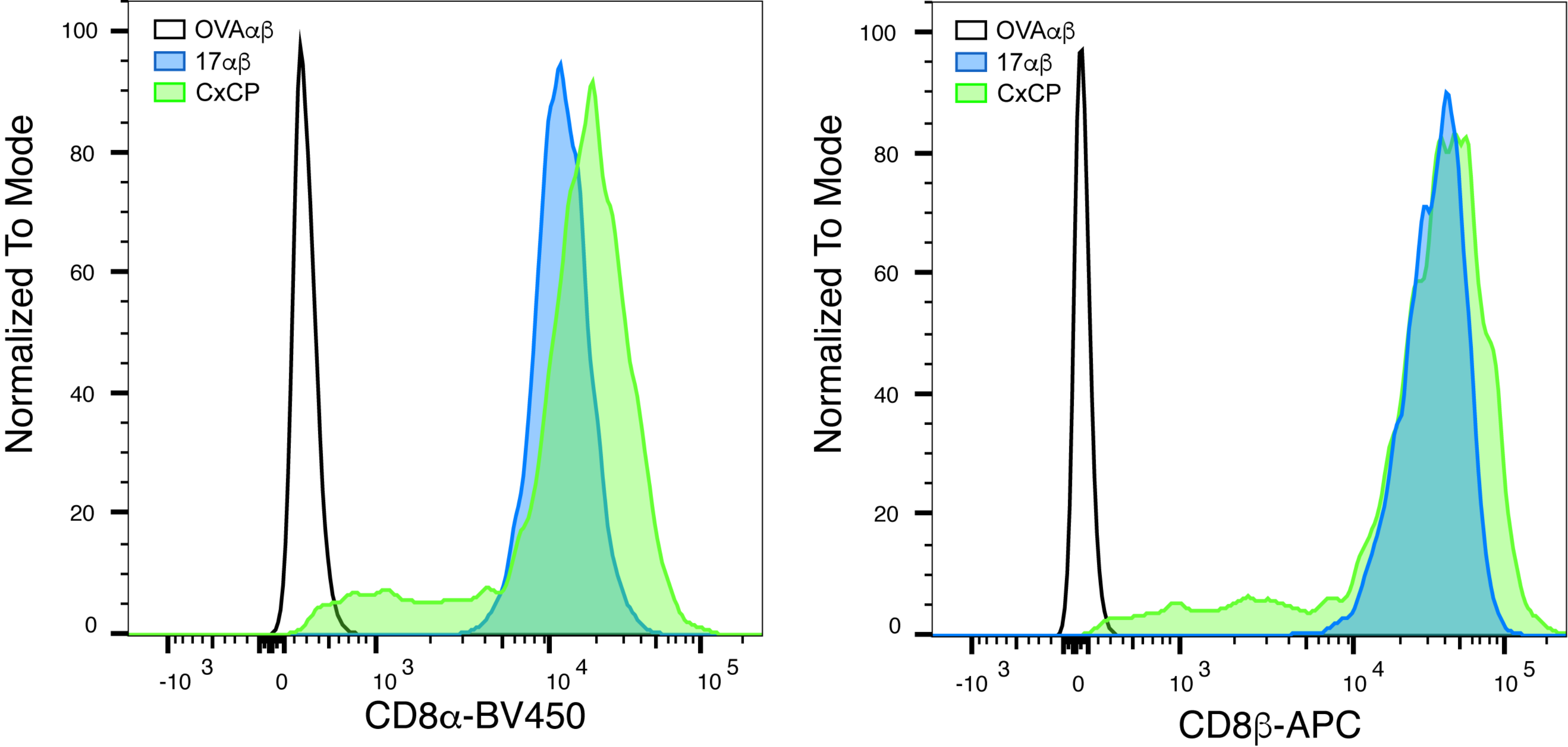

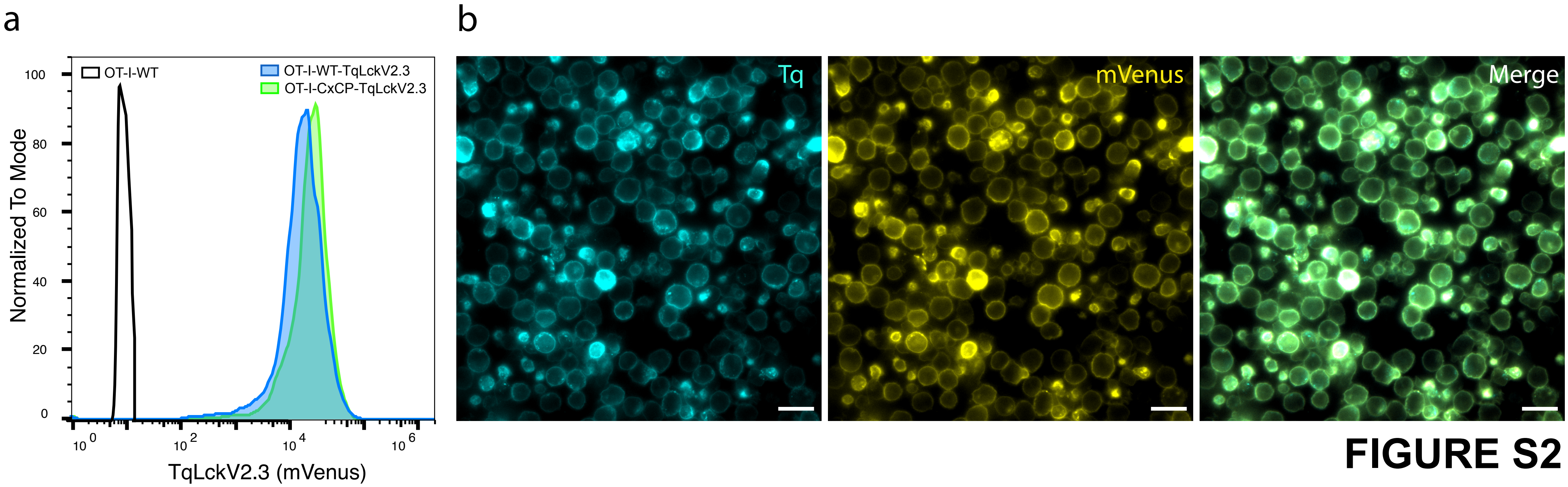

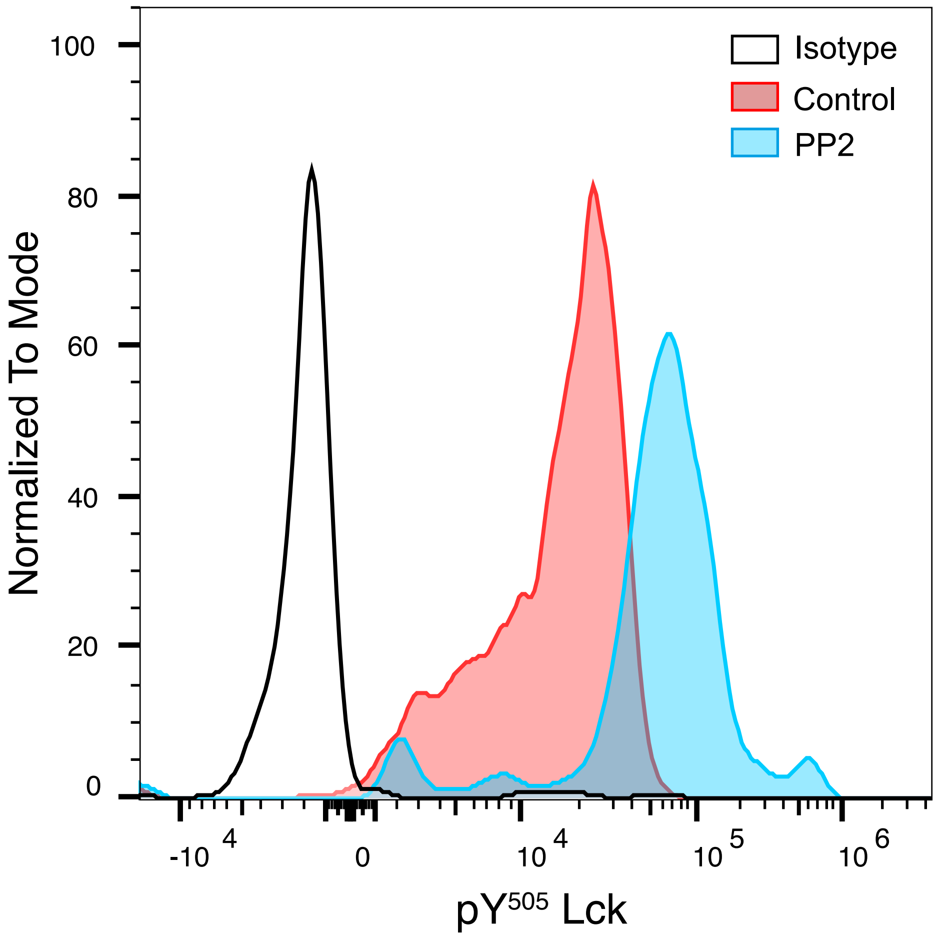

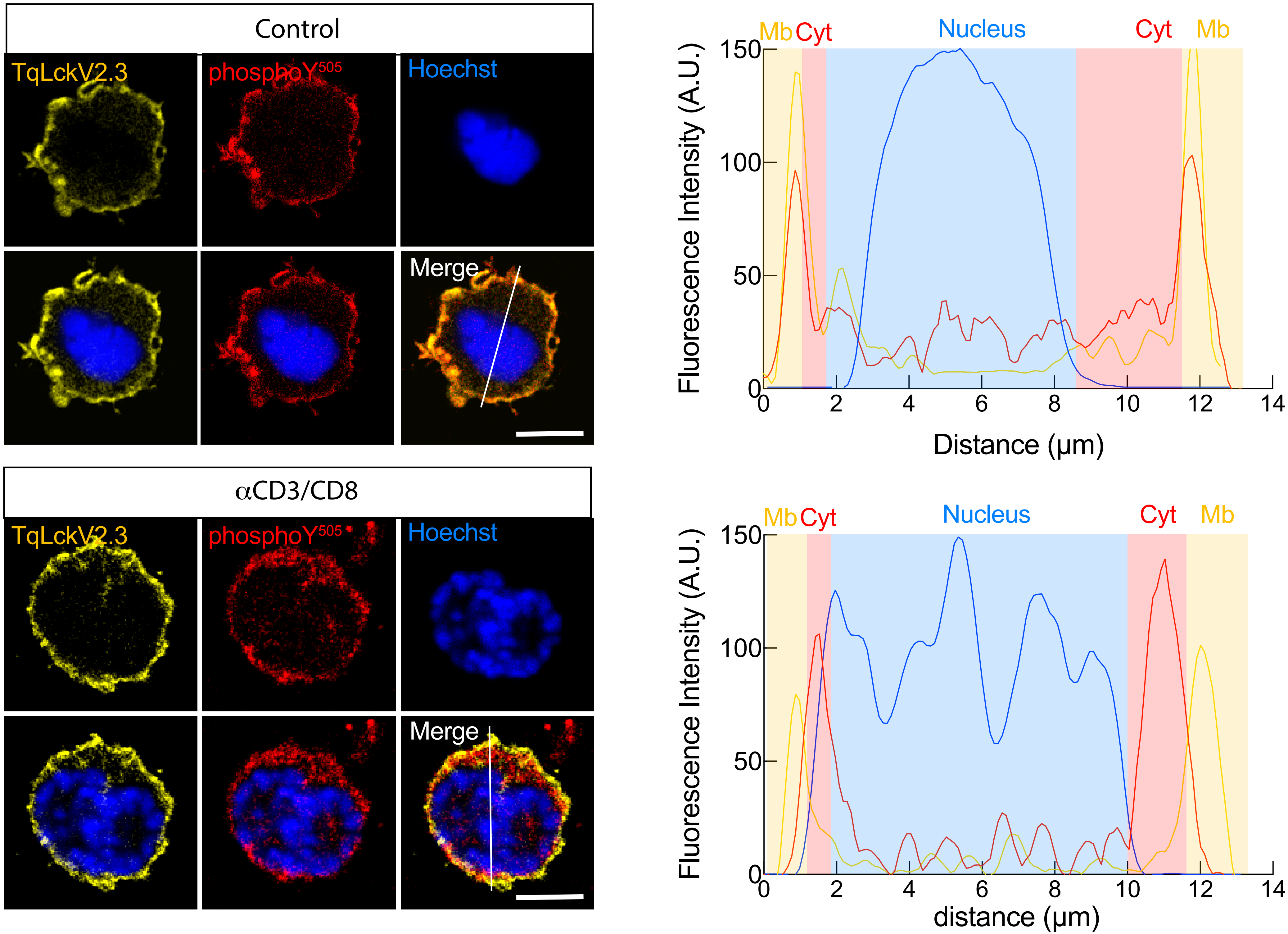
